# Eucalyptus plantations in temperate grasslands: responses of taxonomic and functional diversity of plant communities to environmental changes

**DOI:** 10.1101/2020.10.12.334011

**Authors:** Pamela E. Pairo, Estela E. Rodriguez, M. Isabel Bellocq, Pablo G. Aceñolaza

**Affiliations:** Centro de Investigaciones Científicas y Transferencia de Tecnología a la Producción (CICyTTP-CONICET), Materi y España (3105) Diamante, Entre Ríos, Argentina; Centro Regional de Geomática, Universidad Autónoma de Entre Ríos (CEREGEO-UADER), Ruta 11 km 10.5, Oro Verde (3101) Entre Ríos, Argentina; Laboratorio de Ecología de Comunidades y Macroecología, Departamento de Ecología, Genética y Evolución, IEGEBA, (CONICET-UBA), Facultad de Ciencias Exactas y Naturales, Universidad de Buenos Aires, Ciudad Universitaria, Pabellón 2, Piso 4, CA Buenos Aires (1428), Argentina; Facultad de Ciencias Agropecuarias (FCA-UNER). Ruta 11 Km 10 (3101), Oro Verde, Entre Ríos, Argentina

**Keywords:** functional diversity, plant diversity, grassland, tree plantations, forest cycle

## Abstract

Understanding how human land-uses impact on local communities is required to implement management and conservational policies and practices. Tree plantations have become one of the fastest-growing land uses in recent decades and their impact on biodiversity was evaluated mainly at the taxonomic level. Our aim was to analyze the effects of changes in environmental drivers along the 12 years eucalypt plantations chronosequence on alfa, beta, taxonomic and functional diversity of understory plant communities. We selected nine plantation ages with three replicates per age and three protected grasslands as reference habitat. At each replicate, we established three plots to measure plant species cover and environmental variables, which are expected to change with plantation age. Results showed that species richness and all diversity indices significantly declined with increasing plantation age. Canopy cover, soil pH, and leaf litter were the most important drivers that explained the decline in taxonomic and functional diversity of plants through the forest plantation. Based on the Path analyses results, canopy cover had an indirect relationship with plant functional diversity mediated by leaf litter, soil pH and plant species richness. The results of the association between functional traits and environmental variables have revealed that high dispersal potential, annual, barochorous, and zoochorous plant species were the functional traits more affected by the eucalypt plantations. Given that leaf litter was negatively associated with all diversity facets, we recommend reducing their accumulation within eucalypt plantation to enhance biodiversity conservation and the provision of pampean grassland ecosystem functions.

## 1. INTRODUCTION

The human population is growing more rapidly than ever before and consequently increasing the demand for goods and services provided by ecosystems processes (Sala et al., 2000; Wall and Nielsen, 2012). Among human activities that imply intensive land use, tree plantations with non-native species are expanding worldwide because of the global demand for timber, pulp and biofuel products (FAO, 2015). The effects of land conversion on ecosystem services will largely depend on how native biodiversity responds to the resulting environmental changes. Further studies revealed that that human land uses that preserve vegetation structure and composition of natural biomes are more used by native species than those anthropogenic habitats that cause drastic environmental changes (Filloy et al., 2010; Santoandré et al., 2019). For instance, tree plantations have more structural similarities to natural forests than crop fields, then promoting suitability for forest biodiversity but less than native forest (Calviño-Cancela, 2013). In addition, silvicultural practices and plantation age are also relevant to be considered in studies of persistence of native species (Joelsson et al., 2018).

Several studies have already underlined the negative effects on native species and their related ecosystem functions due to the expansion of forest plantations (Tererai et al., 2013). Besides, the majority of studies on biodiversity changes among different plantation ages have been conducted mainly in forest biomes (Barlow et al., 2007; Jacoboski et al., 2016; Proença et al., 2010; Wu et al., 2015), showing that mature plantations contribute to maintain biodiversity better than young plantations. The opposite pattern was observed for bird and ant communities in tree plantations developed in grassland sites, where the richness and abundance were the lowest in mature plantations and highest in the grasslands (Corbelli et al., 2015; Filloy et al., 2010; Phifer et al., 2016; Santoandré et al., 2019). However, the influence of environmental factors on species biodiversity patterns along the forest cycle remains unresolved.

Eucalypt plantations is one of the most common tree planted forest around the world which have been introduced in pampean grasslands replacing large areas of pastures and crops altering nutrient cycles and abiotic conditions (Phifer et al., 2016; Vega et al., 2009). It has been reported that eucalypt plantations produce nutrient poor litter resulting in low soil nutrient accumulation after decomposition (Ntshuxeko and Ruwanza, 2018). Such changes to soil properties, in turn, affect the structure and composition of soil biological communities, and consequently affects nutrient cycling processes (Kerr and Ruwanza, 2016). Although the allelopathic effects and shading with the increasing canopy cover are the abiotic factors reported why eucalypt plantations reduce understory vegetation and mainly native plant species (Ntshuxeko and Ruwanza, 2018; Zhang et al., 2010), little is known about the link with functional diversity.

Since the beginning of the 21^st^ century, many ecologists have used the functional trait-based approach to complement the traditional taxonomic approach in studies of community richness and composition (Díaz and Cabido, 2001; Petchey and Gaston, 2006). Functional traits are the characteristics of a given organism related to its response to the environment and/or its role in ecosystem functioning (Díaz and Cabido 2001) (e.g. specific leaf area, whether plants are nitrogen-fixing legumes, dispersal mode). Thus, assessing the consequences of environmental disturbances on biological communities and ecosystem functioning through changes in functional diversity (i.e. the value and range of functional trait values present in a community (Díaz and Cabido, 2001)), is critical to identify species traits that can dictate how species respond to environmental changes and to determine their effect on ecosystem function (Lindenmayer et al., 2015; Luck et al., 2012).

Traditional beta diversity was defined as the variation in species composition (Whittaker, 1960), and their study attempt to reveal the assembly mechanisms that drive the variation. More recently, functional beta diversity was defined as the change in ecological functions or species traits between assemblages (Swenson, 2014). The inclusion of both facets of beta diversity is useful for understanding how human activities impact on biodiversity. Vaccaro et al., 2019 have shown that bird composition and functional diversity in cattle pastures were the most similar to native pampean grasslands than other human land uses. However, the effects of tree plantations on the functional diversity of understory vegetation during the forest cycle developing in grasslands are still unclear, even considering that they are among the most threatened and less conserved habitats (Blair et al., 2014). Plants are often used as good biodiversity indicators because they are sensitive to abiotic conditions and land-use changes, are at the base of food webs, and provide habitat for animals influencing animal diversity and distribution (Nic Lughadha et al., 2005).

The study aims to analyze the effects of changes in environmental drivers along the 12 years eucalypt plantations (*Eucalyptus grandis* W. Hill) chronosequence on alfa, beta, taxonomic and functional diversity of understory plant communities. Recognizing the structural and microclimatic changes that occur during the eucalypt forest cycle, we hypothesized that plant taxonomic and functional similarity with the natural habitat (Pampean grasslands) will decline with decreasing environmental similarity between plantation age and grassland sites. Therefore, we expect that 1) as plantation age increases, environmental conditions typical of grasslands will gradually change through the forest cycle (e.g. increase in vegetation stratification, leaf litter depth, canopy cover, and decrease in soil pH); 2) alpha taxonomic and functional diversity will be higher in young plantations than mature plantations; and 3) plant taxonomic and functional similarity (the inverse of beta diversity) with the native grassland will be high in young plantations due to the high environmental similarity. Given that the capability of surviving in disturbed habitats is related with life-history traits (dispersion, establishment, and persistence), our predictions about functional traits are that 1) plant communities in young plantations will have similar functional characteristics to pioneer species e.g. high potential dispersal and shorter lifespan 2) loss of functional traits typical of grassland plant species will occur as plantation age increases and 3) late-successional species may persist at the end of the forest cycle (Marteinsdóttir and Eriksson, 2014; Salgado Negret, 2015).

## 2. MATERIAL AND METHODS

### 2.1. Study area and forest management

The study was conducted in the Entre Ríos Province that includes the northern portion of the Pampa region (the Mesopotamic Pampa subregion), Argentina (Figure 1). Climate is temperate humid, with an average annual temperature of 18° C and annual average precipitation ranging from 1000mm to 1200mm (Morello et al., 2012). The study area presents a rolling relief, well-defined streams bordered by riparian forests (Cabrera, 1971; Rodriguez et al., 2017, 2018). The dominant vegetation type was composed of *Axonopus* sp., *Paspalum* sp., *Digitaria* sp., *Schizachyrium* sp., and *Bothriochloa* sp. (Bilenca and Miñarro 2004). The general history of anthropogenic land use in the region began with extensive ranching during the 20^th^ century, followed by soybean (encouraged by the implementation of direct seeding) and citrus crops in the 21th century (Inta, 2011). Currently, the expansion of eucalypt plantations in the Mesopotamic Pampa is replacing mainly citrus and soybean crop fields, cattle pastures, and native grassland fragments.

**Figure 1.**
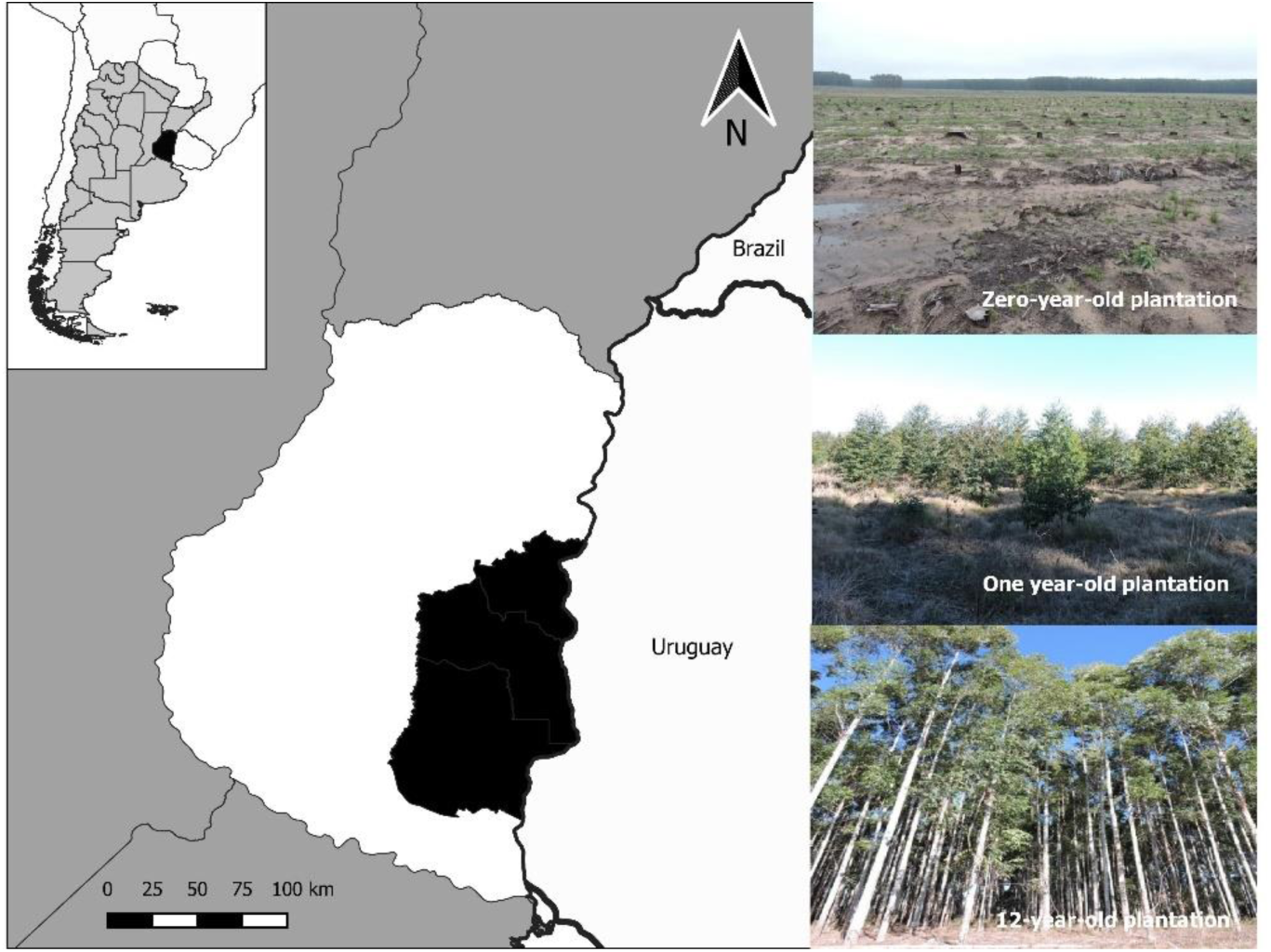
Location of the study area in the pampean grassland in Argentina.

The Pampas region in South America is one of the greatest temperate grassland of the world and harbored one of the fastest growth planted forests in the American continent (Bilenca and Miñarro, 2004). Eucalypt is the primary tree genus planted in the region, promoted by high biomass yield, public incentives, and an emerging market of carbon sequestration (Jobbágy et al., 2006; Phifer et al., 2016). Eucalypt plantations selected in this study were managed with even-aged silviculture, with trees planted 4×2.5 m apart and stocking densities averaging 1000 trees/ha (Aguerre et al., 1995). Herbicide was applied before planting to reduce competition with grasses and herbs. In addition, formicide application was implemented to avoid herbivory from leaf-cutting ants (Aparicio et al., 2005; Vilela et al., 2015). Both treatments were applying at 0-1 year after planting the eucalypt tree. Eucalypt plantations in this area were harvested for pulp and timber 10-12 years, and subjected to silvicultural treatments such as pruning and thinning (Aceñolaza et al., 2013). The first and second pruning and thinning are conducted at 2 years and 7-8 years after plantation establishment, respectively (Aguerre et al., 1995). After the mentioned forest management, harvest residues (leaves, bark, twigs, and branches) are not removed from stands. The forest density was 400-600 trees/ha in mature plantations.

### 2.2. Study design, plant sampling and environmental variables

Eucalypt plantations of nine ages (0, 1, 2, 3, 4, 5, 7, 9, 12 years) were selected to represent the chronosequence; and also established grassland sites to use as reference habitat. The zero-year old plantations referred to the period after planting but less than one year. There were three replicates per plantation age and grasslands for a total of 30 sites. At each site, we established three 16-m^2^ plots (total 300 plots) at least 50m apart from plantation borders to avoid edge effects. During the blooming season from the late spring to early summer (December to January), plant species were identified at each plot, assigned percentage cover and measured environmental and dasometric variables related with changes that occur during the forest cycle. The collected specimens were identified to species or morphospecies level consulting the main territorial floras (Burkart, 1969, 1974, 1979, 1987). Moreover, canopy cover of eucalypt trees was estimated using five digital photos taken from 1.5 m above the ground toward the canopy at each site. The percent canopy on each photo was analyzed with ImageJ as percentage of pixels with vegetation (Peyras et al., 2013). The average of leaf litter depth (mostly composed by eucalypt leaves) was estimated by measuring the 10 random points per site. Similarly, average of tree height and diameter at breast height (DBH) were measured using 10 eucalypt trees. Finally, 10 soil subsamples at 0–20 cm depth were taken by walking a ‘zig-zag’ pattern from each site and mixed for homogenization. From each soil sample, total soil carbon, nitrogen, phosphorus (Bray& Kurtz), and soil pH were determined in the laboratory.

### 2.3 Taxonomic diversity

To estimate alpha taxonomic diversity, we counted plant species richness per site. Moreover, to estimate beta taxonomic diversity, we calculated the Sorensen dissimilarity index (β_sor_) between each plantation age, and the native grassland pooled the three grassland sites to better represent the regional pool of species.

### 2.4 Functional diversity

#### 2.4.1 Selection of functional traits

Plant species that occurred once were removed from total species recorded for measuring functional diversity indices. The main reason is that dominant plant traits are known to strongly influence ecosystem processes, indicating how the community in general responds to the environment (Diaz et al., 1998). In general, plant communities have a typical structure with a relatively small number of dominant species and a large number of subordinate or minor species that account for a low proportion of the biomass (Grime, 1998; Carreño-Rocabado et al., 2015). Plant species recorded on the plots were scored for four functional traits associated with community responses to disturbances: growth form, dispersal syndromes, dispersal potential and life history (see Table 1) (Pérez-Harguindeguy et al., 2013).

**Table 1.** Description of functional traits selected of plant species in eucalypt plantations of the pampean grasslands. Information were collected from (A. Burkart, 1969; Dawson et al., 2017; Pérez-Harguindeguy et al., 2013; Rodriguez et al., 2018).

| Functional<br>Trait | Values | Description |
| --- | --- | --- |
| Growth form | Tree, herb, subshrub | Associated with<br>ecophysiological<br>adaptation. |
| Dispersal<br>syndromes | Anemochory,<br>anemozoochory, barochory,<br>zoochory | Associated with the<br>covered distances,<br>the routes and the<br>destination. |
| Dispersal<br>potential | High, medium, low | Amount of seeds<br>produce by one<br>individual. The<br>capacity to survive |
|  |  | in disturbed |
|  |  | habitats is often |
|  |  | correlated with a |
|  |  | high dispersal |
|  |  | potential. |
| Life history | Annual, biennial, perennial | Indicator of |
|  |  | population |
|  |  | persistence, adult |
|  |  | plant longevity |

#### 2.4.2 Functional diversity indices

We calculated functional dispersion (FDis) based on previous functional traits selected using the FD package (Laliberté and Legendre, 2010). FDis is defined as the mean distance in a multidimensional trait space of individual species to the centroid of all species, weighted by their relative abundances. To estimate beta functional diversity along eucalypt chronosequences (considering grassland sites as the reference habitat), we used trait matrix to build a dendrogram tree using the Jaccard index because functional traits are categorical variables and to avoid that not sharing traits increase the similarity between two species (Legendre and Legendre, 2012). Then, both dendrogram tree and presence/absence data per site were used to calculate the Fsor index using the PICANTE package (Kembel et al., 2018), which is analogous to a traditional Sorensen’s Index. Fsor represents the proportion of dendrogram branch lengths shared by two assemblages (Swenson et al., 2011).

### 2.5. Data analysis

To describe how environmental variables changed with plantation age, we carried out Principal Component Analysis (PCA). To test changes in species richness (alpha taxonomic diversity) along the eucalypt forest cycle, we performed Generalized Linear Models (GLM) with Negative Binomial distribution because we had a count variable (species richness) and to reduce overdispersion. We modeled similarity (1-β_SOR_) with beta distribution and performed beta regression models to explain the variability along the eucalypt forest cycle. Beta regression models are based on the assumption that the response variable is beta-distributed and it was developed for situations where the dependent variable can take values between 0-1 as the case of the Sorensen index (Cribari-Neto and Zeileis, 2010).The results of beta diversity are presented in terms of similarity in composition between communities (1-β_SOR_). To identify the main environmental variables explaining changes of taxonomic diversity along the eucalypt forest cycle, we used a model-selection approach with Akaike Criterion (AICc), as we handled the corrected version for small sample sizes (Burnham and Anderson, 2002). Models with ΔAICc < 2 were considered as equivalent to the minimum AICc model and hence more robust to explain the observed variability (Zuur et al., 2010). Goodness-of-fit was evaluated by examining plots of standardized residuals vs. predicted and checked normal distribution. It is important to remark that environmental similarity indices (1-Gower index) between each environmental variable respect to grassland sites were calculated for beta taxonomic analyses. To assess changes in alpha and beta functional diversity along the eucalypt forest cycle, we modeled FDis and Fsor with Normal Distribution. As noted previously for taxonomic diversity analyses, the model selection approach was implemented to identify the environmental variables that explained the variability in alpha and beta diversity along the eucalypt forest cycle.

Structural equation models (SEMs) are a statistical technique that unite multiple variables in a single causal network, thereby allowing simultaneous tests of multiples hypotheses (Grace, 2006). SEMs test the direct and indirect effects on pre-assumed causal relationships (Fan et al., 2016). Piecewise SEM expands upon traditional SEMs by introducing a flexible mathematical framework that can incorporate a non-normal distributions, hierarchical structures and different estimation procedures (Lefcheck, 2016). We then performed a piecewise SEM to analyze the direct and indirect effects on the alfa functional diversity (FDis), including the environmental variables with the lowest AICc from the model selection and the plant species richness. Some researchers recommend a minimum sample size of five cases per free parameter in the model resulting in six possible relationships to be tested (Fan et al., 2016). The direct relationship between species richness and alpha functional diversity was modeled with Poisson distribution. Fisher’s C statistic to a chi-square distribution was calculated to evaluate the global goodness-of-fit of the model using the ‘psem’ function from the ‘piecewiseSEM’ package (Lefcheck et al., 2019).

In order to identify associations between functional traits and environmental variables, we used fourth-corner analysis, i.e. measured through the interaction terms between each environmental variable and trait variables (Brown et al., 2014). The sign and magnitude of the interaction coefficients denoted the nature and strength of the association of the trait–environment relationship, respectively. We fitted a linear regression model, using all functional traits and environmental variables and the LASSO penalty was estimated via cross-validation. LASSO is a method of penalized likelihood which imposes a constraint on estimates of model parameters (Hastie et al., 2009).

All statistical analyses were conducted in R 3.2.0 (R Core Team 2015). Beta regression was performed using functions in the R package ‘betareg’ (Cribari-Neto and Zeileis, 2010), and candidate models were selected with the R package ‘MuMIn’ (Barton, 2018).

## 3. RESULTS

We recorded a total of 247 plant species/morphospecies along the eucalypt forest cycle and grassland sites in the Mesopotamic Pampa (30 sites in total). We identified 47 species/morphospecies in grassland sites, 32 species/morphospecies in both grassland and eucalypt plantations and 168 species/morphospecies only in eucalypt plantations (Appendix A). Moreover, we identified 49 families and the most dominant were Poacea (47 spp.), Compositae (47 spp.) and Leguminosae (16 spp.).

Regarding environmental changes, the first two axes of PCA analyses explained 87% of the variation (Figure 2). PCA axis 1 arranged sites according to eucalypt growth from negative to positive values. Grassland sites, 0-year-old and 1-year-old plantations have similar environmental conditions: high levels of soil pH, phosphorus content and temperature. Moreover, the rest of the plantations were high canopy cover, DBH and leaf litter depth.

**Figure 2.**
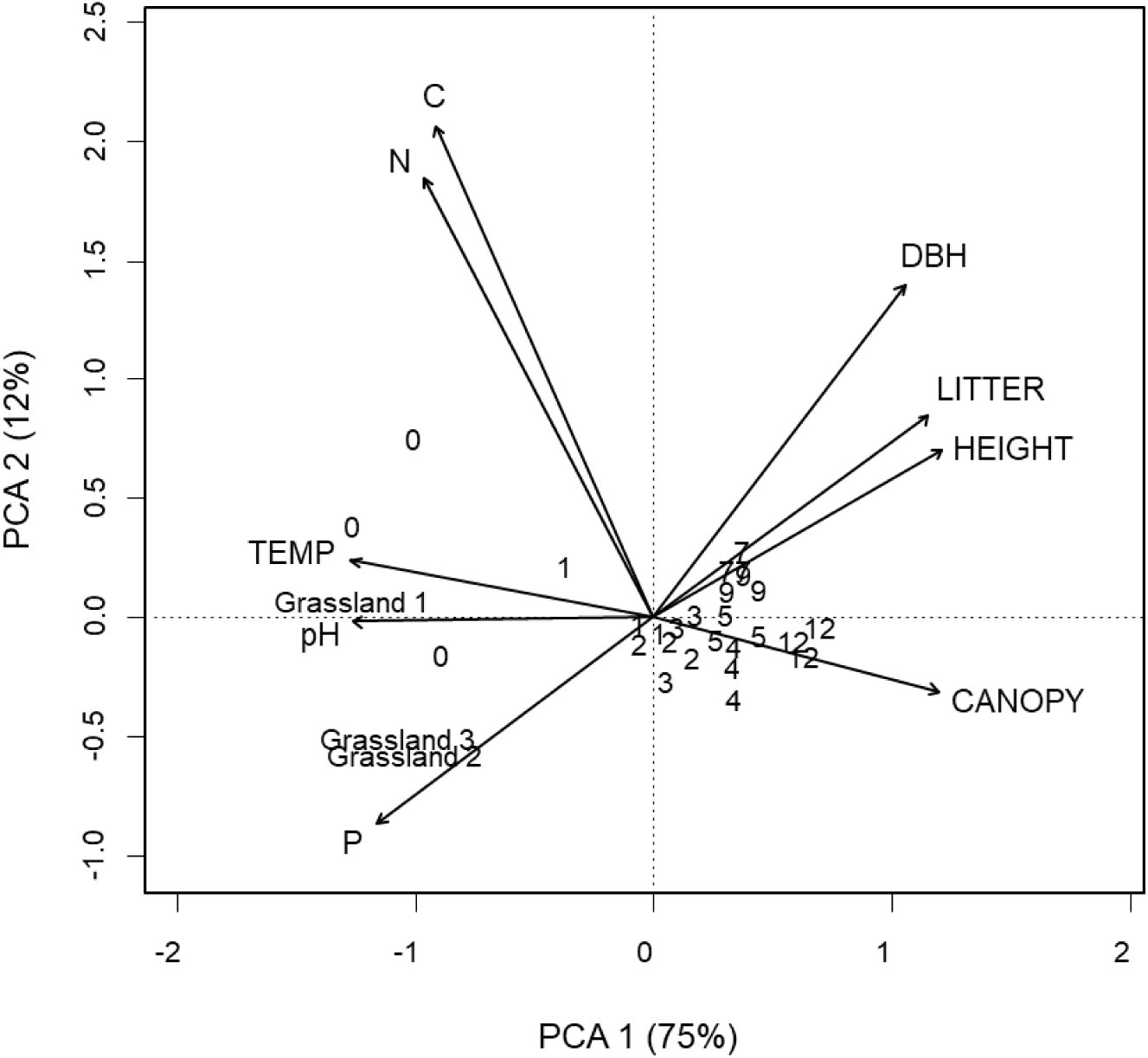
Biplot from principal component analyses (PCA) of the environmental variables measured along eucalypt forest cycles and grassland sites of the Mesopotamic Pampa. The first and second axes of the PCA are shown. The percentage of variance explained by each axis is given into brackets. CANOPY= canopy cover, LITTER= leaf litter depth, HEIGHT= eucalypt tree height, DBH= diameter at breast height, P= phosphorus content in soil, C= carbon content in soil, N=nitrogen content in soil. Numbers represent the plantation ages.

Plant species richness and taxonomic similarity significantly decreased as plantation age increased (Z= -3.49, p < 0.001; Z=-2.689, p= 0.007, respectively) (Figure 3 A, B). In accordance, the observed alpha functional diversity and functional similarity declined as plantation age increased (T= -2.58, p= 0.02; Z=-2.74, p= 0.01, respectively) (Figure 3 C, D).

**Figure 3.**
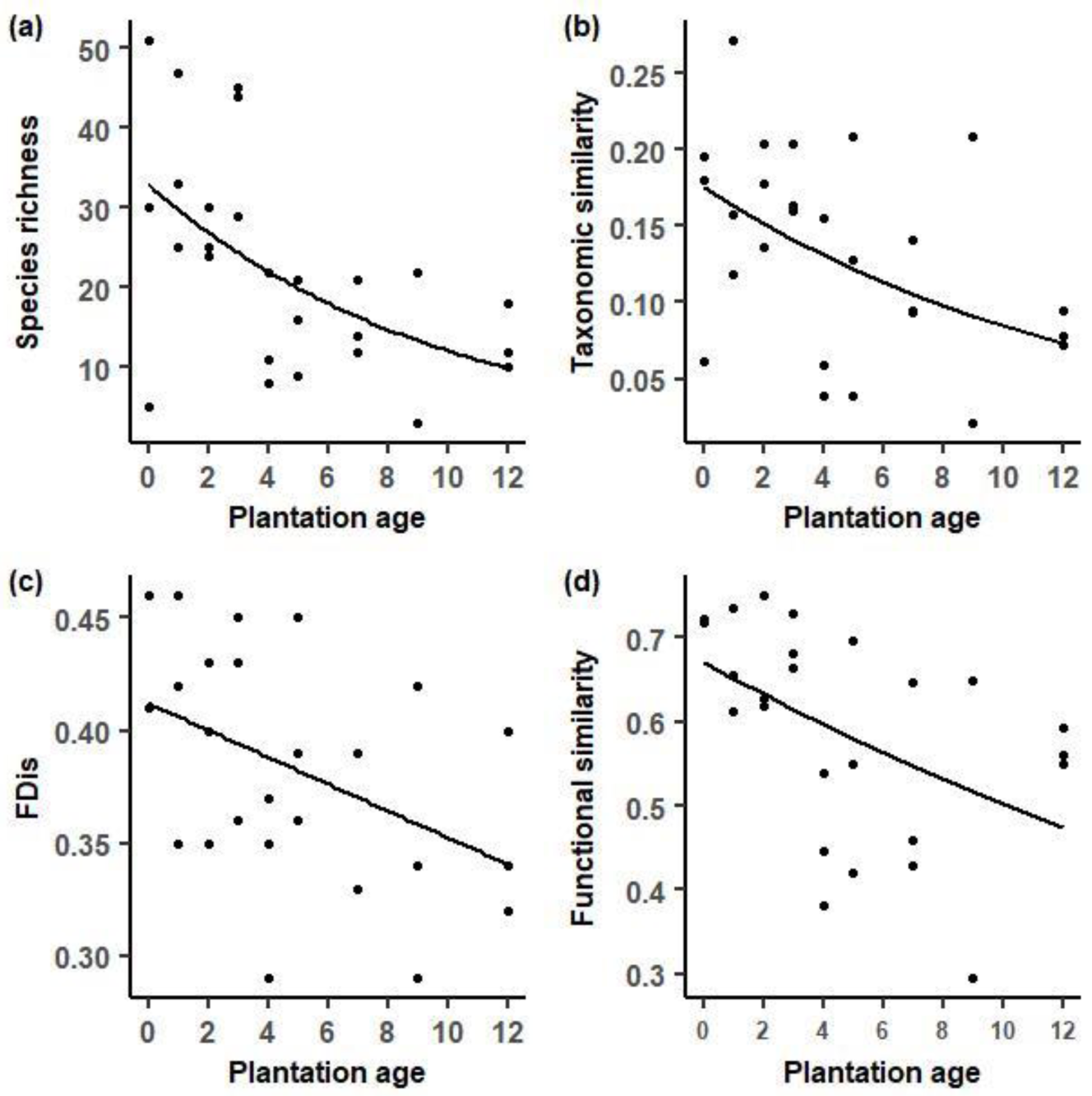
Changes of plant diversity facets along eucalypt forest cycle in grassland context. a) Species richness (alpha taxonomic diversity), b) taxonomic similarity (1-βSor) c) alpha functional diversity (FDis) d) functional similarity (1-FSor). Grassland sites were pooled for measuring similarity indexes.

Our results showed that plant species richness was negatively affected by soil pH, leaf litter depth, and nitrogen content (Table 2). In addition, we found that overall taxonomic similarity was also explained by similarity in leaf litter depth and canopy cover between grassland sites and the eucalypt forest plantations. The decline of alpha functional diversity was affected by the increase of canopy cover and leaf litter depth. Finally, the decline in functional similarity along eucalypt forest cycle was explained by the increase of similarity in leaf litter depth, soil pH, and nitrogen content. The models with ΔAICc > 2 have been present in Appendix B, C, D, and E.

**Table 2.** Summary of best models explaining variation of diversity indices in eucalypt chronosequence in the Mesopotamic Pampa. Best models (with ΔAICc < 2) are presented as these models are typically considered to present similar statistical support. d.f.: degree of freedom, AICc, %E.V. = percent of explained variability of each model; pseudo R^2^ was used for beta taxonomic and functional diversity, and adjusted R^2^ for alpha functional diversity. For GLM with negative binomial distribution, we calculated the explained variability of each model as the ratio: (null deviance-residual deviance)/null deviance. sN= similarity in nitrogen content, sCanopy=similarity in canopy cover, sLitter=similarity in leaf litter depth.

| Diversity | N/sN | Canopy/ sCanopy | Soil pH | Litter/ sLitter | d.f. | AICc | %E.V |
| --- | --- | --- | --- | --- | --- | --- | --- |
| Alpha Taxonomic | - | - | -0.48 | -0.15 | 4 | 207.17 | 41 |
|  | 2.68 | - | -0.59 | -0.15 | 5 | 208.07 | 45 |
|  | - | - | - | -0.12 | 4 | 209.07 | 30 |
| Taxonomic similarity | - | - | - | 0.75 | 3 | -68.27 | 15 |
|  | - | -0.47 | - | 1.02 | 4 | -67.31 | 18 |
| Alpha functional | - | -0.001 | - | - | 3 | -87.37 | 20 |
|  | - | -0.001 | - | -0.003 | 4 | -85.93 | 22 |
| Functional similarity | - | - | - | 0.41 | 3 | -34.40 | 22 |
|  | 0.29 | - | -0.56 | 0.35 | 5 | -33.27 | 41 |
|  | - | - | -0.33 | 0.35 | 4 | -33.19 | 29 |
|  | 0.15 | - | - | 0.41 | 4 | -32.55 | 26 |

Piecewise SEM analysis revealed that the plant species richness affected positive and directly on alpha functional diversity (Fig. 4). In addition, our model showed that the effect of the canopy cover on plant species richness is mediated by leaf litter depth and soil pH. Leaf litter depth had a direct and indirect influence on plant species richness. The analysis reproduced the data well based on a comparison of the Fisher’s C statistic to a chi-square distribution (C_8_= 5.56, p = 0.7).

**Figure 4.**
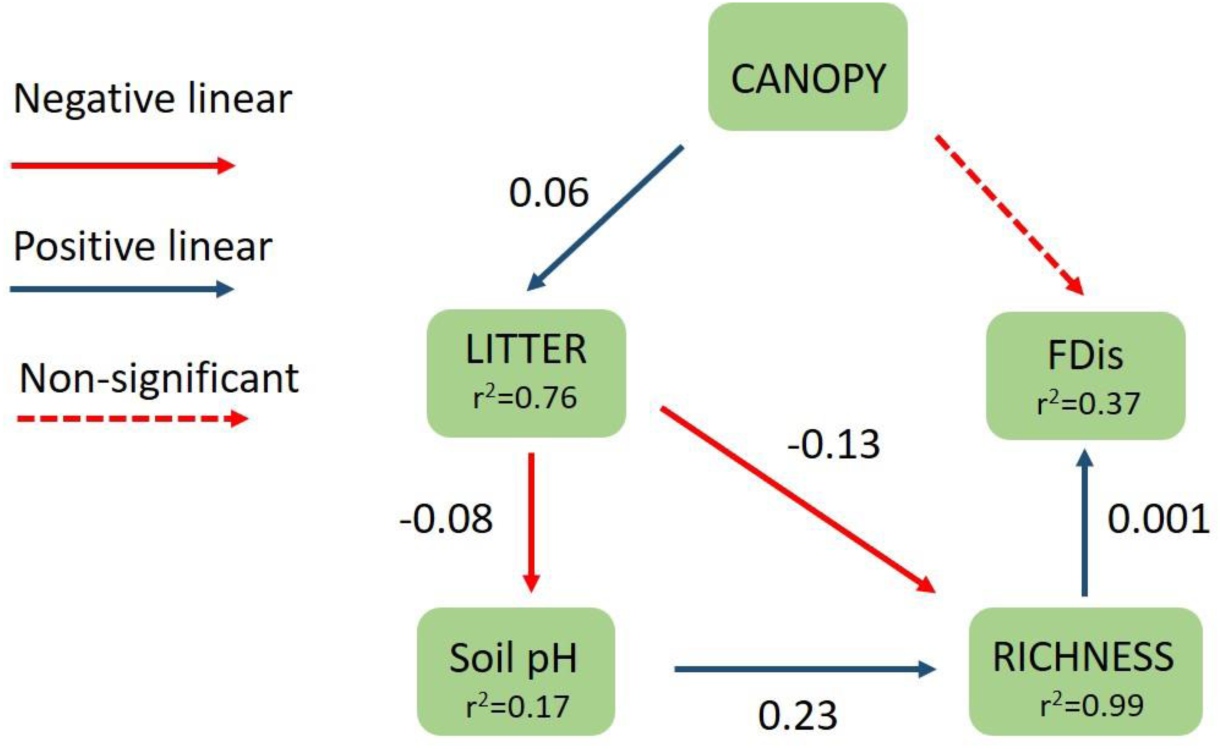
Piecewise SEM model showing the direct and indirect effect of CANOPY (canopy cover), LITTER (leaf litter depth), soil pH and RICHNESS (plant species richness) on alpha functional diversity (FDis). Arrows represent unidirectional relationships among variables and their standardized coefficients are shown (p < 0.05). Dashed arrow represents nonsignificant relationship (p > 0.05). R^2^ for component models are given in the boxes of response variables.

Results of the fourth corner analysis revealed a positive and strong association between canopy cover and anemozoochory as dispersion syndrome and, conversely, a negative and strong association with barochorous species (Fig. 5). In addition, leaf litter depth had a negative and strong association with plants with high dispersal potential, perennial plants and zoochory and barochory as dispersal syndromes. Soil pH had a negative association with perennial plants and herbs whereas N had a negative and strong association with annual plants and anemozoochory. Finally, nitrogen content in soil had a negative strong association with annual plants and anemozoochory as dispersal syndrome.

**Figure 5.**
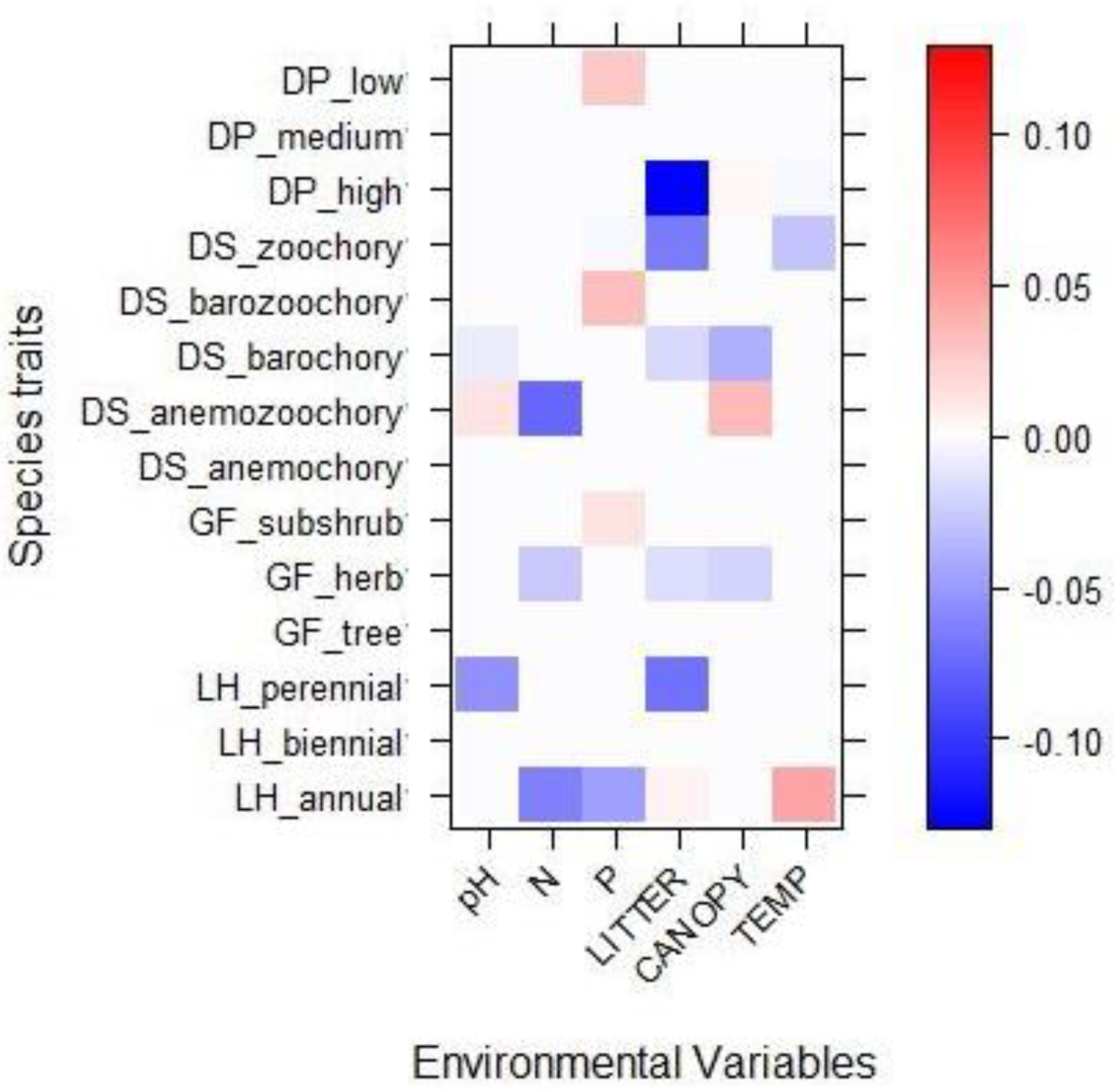
Results of fourth-corner modelling for plant trait interactions with environmental changes that occur along eucalypt forest cycle in the Mesopotamic Pampa. Red represents a positive association and blue a negative association. Color grading show the direction and strengths of standardized coefficients of fourth-corner models for all environment/trait interaction terms from GLM-LASSO modelling. LH= life history, GF= growth form, DS= dispersal syndrome, DP= dispersal potential. Acronyms for environmental variables are shown in Figure 2.

## 4. DISCUSSION

Our findings reveal that changes in soil pH, canopy cover, nitrogen content in soil, and leaf litter from native grassland to mature plantations were the major drivers of the decline in the taxonomic and functional diversity of plants. To our knowledge, this is the first time to simultaneously compare plant taxonomic and functional diversities and their relationship with environmental variables providing new insights into the effects of plantation cycle on grassland plant communities. Furthermore, we propose a model that includes soil (soil pH) and structural variables (litter depth, canopy cover) to explain the variation in plant taxonomic and functional diversity along the eucalypt forest cycle. Eucalypt tree growth imposed novel abiotic conditions to pampean grasslands could modify resource availability to plant species (light, water, and nutrients) (Barbier et al., 2008).

As expected, alpha taxonomic and functional diversity decreased with plantation age, and it was explained by the changes in soil pH, leaf litter depth, nitrogen content in soil and canopy cover along the eucalypt forest cycle. Similarly, decreasing in understory plant species in the first four years after eucalypt plantation establishment was found within the Danling region in China where the landscape mosaic is composed of commercial plantations, cultivated land and unmanaged forests (Zhang et al., 2014). The accumulation of leaf litter has been considered a factor of allelopathy by eucalypt plantations as a response to competition, through a reduction in seed germination and growth of native plants (Zhang and Fu, 2009). Certain phenolic acids and volatile oils are released from the leaves, bark and roots of some *Eucalyptus* spp. act as allelopathic agents and are harmful to other plant species (Florentine and Fox, 2003). Jobbágy et al., (2006) found that eucalypt plantations in grasslands acidify the soil mainly as a result of the release of phenolic acids. Alternatively, soil acidification could be caused by an allelopathic response (Souto et al., 2001) and would difficult the availability of nutrients for understory plants. On the other hand, canopy cover was a relevant factor in decreasing plant functional diversity and consequently impact on functions that plant species performed. Furthermore, light availability on the forest floor, microclimatic conditions (i.e temperature and humidity) and rainfall patterns are closely dependent on canopy cover (Suggitt et al., 2011; Zellweger et al., 2017). The forest canopy generate a rainfall partitioning into interception, throughfall and stemflow and then affect soil moisture patterns, infiltration, groundwater recharge and water yield (Silveira and Alonso, 2009). Considering that high nitrogen availability in the soil could stimulate microbial activity and then an increase of decomposition rate (Mendoza et al., 2017), the positive association between plant species richness and nitrogen content in soil would be related to the high microbial activity allowing that certain soil nutrients are available. Therefore, the eucalypt forest cycle modified habitat characteristics limiting the establishment of some plant species and their functional traits because of their ecological niches were not compatible with the new environmental conditions.

Regarding taxonomic and functional similarity results, canopy cover, leaf litter depth, soil pH, and nitrogen content in soil were the most important factors which negatively affect grassland plant species. As we have reported previously, leaf litter could produce soil acidification and then allelopathic effects on plant species and would avoid that typical grassland species could remain. The presence of some plant species in eucalypt plantation and their absence in grassland sites (Appendix A) could be explained by the dispersion of some species from riparian forests of the Uruguay River (Rodriguez et al., 2017) to mature plantations. For example, we have recorded the presence of *Blepharocalyx salicifolius, Teucrium vesicarium*, and *Eugenia myrcianthes* in mature plantations, whose dispersion would be favored by high similarity in habitat structure and microclimatic conditions with riparian forest (Corbelli et al., 2015). Besides, studies have reported that tree plantations are hot-spots of plant invasion and threaten the remnants of semi-natural vegetation (Csecserits et al., 2016; Piwczynski et al., 2016). In addition, studies that evaluated changes in plant species composition developed in landscapes dominated by forest have shown the species composition changes considerably due to the increasing richness of invasive species in commercial plantation (Amazonas et al., 2018; Grass et al., 2015; Jin et al., 2016; Verstraeten et al., 2013; Wu et al., 2015). On the other hand, other important factors that could influence the variability in functional diversity are management practices carried out in eucalypt plantations of pampean grasslands. Particularly forest management that invokes increased light availability through the opening spaces, for instance thinning, may effectively increase overall plant species and so biodiversity at different scales (Haughian and Frego, 2016; Piwczyński et al., 2016; Smith et al., 2007). Nevertheless, the authors have remarked that this increase would be consistent with the increase of invasive plant species within tree plantations, then the original native ecosystem functions may have been damaged. Our results suggested that thinning (in 2-year and 7-year eucalypt plantation) not increase plant diversity due to all diversity indexes showed a marked decrease in their values.

The proposed model explaining the effect of the eucalypt forest cycle on alpha functional diversity indicated that canopy cover had an indirect relationship mediated by leaf litter, soil pH, and species richness. Furthermore, the model has shown that the species richness decline was linked with the loss of unique functional traits suggesting low functional redundancy in grassland plant communities of Mesopotamic Pampa. The increase of leaf litter depth had a direct and indirect relationship with species richness due likely to the soil acidification and physical restrictions for plant growth. Based on the coefficients of the model, the impact caused by the physical restriction on species richness (and functional diversity) would be higher than the soil acidification and the allelopathic effects associated. Future researches may incorporate other abiotic factors such as nitrogen in the soil available to plants and humidity, to disentangle the effects of environmental changes on functional diversity.

Results of the associations between life-history traits and environmental variables showed that high dispersal potential, annual, barochorous and zoochorous plant species were the functional traits further affected by the growth of eucalypt plantations. Furthermore, leaf litter, canopy cover, pH and nitrogen content in soil were the environmental variables with a large impact on life-history traits that could act as ecological filters promoting that only species with certain combinations of life traits may pass through the filters. In addition, plant species that occurred in young plantations had similar characteristics to fast-growing pioneer species common in early-successional stages: high amount of seed production, annual lifeform that support our hypothesis. This concurs with the general pattern that greater dispersal ability allows quicker response following disturbance (Connell and Slatyer, 1977). Moreover, the increase of leaf litter depth could be a filter for grassland species which are characterized by high dispersal potential. On the other hand, zoochory and barozoochory were the dispersal syndromes affected by canopy cover and leaf litter that may impact on ecosystem services. However, we expected that anemochorous and anemozoochorus species would have been presented in young plantations not only for their high dispersal abilities but also are characteristics of grassland species which can tolerate the environmental conditions because of the similarity with grassland habitats. Our results are consistent with other studies that found plant traits of life history were associated positively with disturbance intensity (Pedley and Dolman, 2014). Assemblages after disturbance comprised few plants with wind-dispersed seed, consistent with selection for species with better dispersal ability. The observed trait composition could be the response to high disturbance intensity (i.e formicide and glyphosate application) in the early years of eucalypt plantation growth. This set of traits was more likely to colonize new environments and have better abilities to find the necessary resources (Duncan et al. 2003). Changes in trait composition towards generalist species in response to land-use intensification have also been reported for other traits and arthropod taxa in grasslands of Germany (Birkhofer et al., 2015; Mangels et al., 2017). At the end of the eucalypt forest cycle, the plant species presented traits related to late-successional species (less dispersal abilities, perennial lifeform) and differential dispersal syndrome with respect to young plantations. It is important to notice that successional processes are driving plant community changes, not only the environmental changes caused by the growth of eucalypt plantations, and our study design cannot separate both effects.

## CONCLUSION

Assessing the changes in taxonomic and functional diversity of plant species along the eucalypt forest cycle highlighted the importance of integrating approaches. Our results demonstrated that facets and components of grassland plant diversity were negatively affected by novel environmental conditions provided by eucalypt plantation growth. Leaf litter depth was the most important driver in plant diversity decline may cause loss of native species in pampean grasslands and consequently could affect resource availability for herbivores, species interactions, trophic networks and ecosystem functioning. Our findings could be useful in places where eucalypt plantation is expanding within open habitats, especially in South America and Africa. We recommend to regional stakeholders that conservation efforts should be addressed to reduce litter accumulation within eucalypt plantations. Finally, our approaches and results may contribute to develop practical management guidelines aimed at enhancing the value of plantations for biodiversity conservation.

## Supporting information

appendixA

appendixB

appendixC

appendixD

appendixE

## ACKNOWLEDGEMENTS

We dedicate this paper, *in memoriam*, to our dear colleague and friend M. Isabel Bellocq who passed away during the writting process. She will continue to be our inspiration as a sweet and kind person and excellent community ecologist. We thank Carolina Pinto and Raúl D’Angelo for their assistance in fieldwork. We are very grateful to the staff of Reserva “El Potrero”, “Estancia El Palmar”, “Beyga SA”, “Aserradero Ubajay”, “Distrimader” and El Palmar National Park for allowing us to work on their properties. This study was supported by Unidad para el Cambio Rural (PIA 14006) and Consejo Nacional de Investigaciones Científicas y Técnicas (CONICET).

