## appendixA for "Eucalyptus plantations in temperate grasslands: responses of taxonomic and functional diversity of plant communities to environmental changes"

**Appendix A.** List of plant species/morphospecies recorded in grassland sites and eucalypt plantation in Mesopotamic Pampa of Entre Rios Province, Argentina. Nomenclatural and author data of plants follow The Plant List (The Plant List 2013).

| **Family** | **Plant species** | **Grassland** | **Forest plantation** |
| --- | --- | --- | --- |
| Amaranthaceae | *Amaranthus sp.* |  | x |
|  | *Amaranthus hybridus subsp. quitensis* (Kunth) Costea & Carretero |  | x |
|  | *Amaranthus viridis L.* |  | x |
|  | *Froelichia interrupta* (L.) Moq. | x |  |
|  | *Iresine diffusa* Humb. & Bonpl. ex Willd*.* |  | x |
|  | *Pfaffia sp.* |  | x |
|  | *Pfaffia gnaphalioides* (L.f.) Mart. |  | x |
| Amaryllidaceae | *Nothoscordum sp.* |  | x |
| Apiaceae | *Cyclospermum leptophyllum (*Pers.) Sprague |  | x |
|  | *Daucus pusillus* Michx. | x | x |
|  | *Eryngium horridum* Malme. |  | x |
|  | *Eryngium nudicaule* Lam. |  | x |
|  | *Eryngium sp.* |  | x |
|  | *Eryngium elegans* Cham. & Schltdl*.* |  | x |
| Apocynaceae | *Mandevilla petraea* (A.St.-Hil.) Pichon | x |  |
|  | *Asclepias curassavica* L. |  | x |
|  | *Morrenia brachystephana* Griseb. |  | x |
|  | *Oxypetalum solanoides* Hook. & Arn. |  | x |
| Araliaceae | *Hydrocotyle sp.* |  | x |
| Arecaceae | *Butia yatay* (Mart.) Becc. |  | x |
| Aristolochiaceae | *Aristolochia sp.* |  | x |
| Boraginaceae | *Echium plantagineum* L. |  | x |
|  | *Heliotropium amplexicaule* Vahl |  | x |
|  | *Heliotropium nicotianifolium* Poir. |  | x |
| Cactaceae | *Opuntia sp.* | x |  |
| Calyceraceae | *Acicarpha tribuloides* Juss. | x | x |
| Campanulaceae | *Triodanis perfoliata subsp. biflora* (Ruiz & Pav.) Lammers |  | x |
|  | *Wahlenbergia linarioides* (Lam.) A. DC | x | x |
| Caryophyllaceae | *Paronychia communis* Cambess. |  | x |
|  | *Silene antirrhina* L. | x |  |
|  | *Silene sp.* | x |  |
|  | *Spergula sp.* | x |  |
|  | *Stellaria media* (L.) Vill. |  | x |
| Commelinaceae | *Commelina diffusa* Burm.f. |  | x |
|  | *Commelina erecta* L. | x | x |
|  | *Commelina platyphylla* Klotzsch ex Seub. |  | x |
|  | *Commelina sp.* | x |  |
|  | *Tripogandra glandulosa* (Seub.) Rohweder | x | x |
| Compositae | *Acanthostyles buniifolius* (Hook. ex Hook. & Arn.) R.M.King & H. Rob. |  | x |
|  | *Achyrocline satureioides* (Lam.) DC*.* | x | x |
|  | *Acmella grisea* (Chodat) R.K.Jansen |  | x |
|  | *Ambrosia tenuifolia* Spreng. |  | x |
|  | *Aster squamatus* (Spreng.) Hieron*.* |  | x |
|  | *Baccharis articulata* (Lam.) Pers. |  | x |
|  | *Baccharis dracunculifolia* DC. | x | x |
|  | *Baccharis pingraea* DC. |  | x |
|  | *Baccharis* *rufescens* var. *rufescens* | x |  |
|  | *Baccharis salicina* Torr. & A.Gray |  | x |
|  | *Baccharis sp.* | x | x |
|  | *Baccharis sp.1* |  | x |
|  | *Baccharis trimera* (Less.) DC. |  | x |
|  | *Bidens sp.* |  | x |
|  | *Calea uniflora* Less. | x |  |
|  | *Campuloclinium macrocephalum* (Less.) DC*.* |  | x |
|  | *Carduus acanthoides* L. |  | x |
|  | *Cirsium vulgare (Savi)* Ten. |  | x |
|  | *Erigeron bonariensis L.* | x | x |
|  | *Eupatorium inulifolium* Kunth |  | x |
|  | *Eupatorium sp.* | x |  |
|  | *Eupatorium sp. 1* |  | x |
|  | *Gnaphalium americanum* Mill. |  | x |
|  | *Gamochaeta argentina* Cabrera | x | x |
|  | *Gamochaeta simplicicaulis* (Willd. ex Spreng.) *Cabrera* | x | x |
|  | *Gamochaeta stagnalis* (I.M.Johnst.) Anderb. |  | x |
|  | *Gnaphalium filagineum* DC*.* |  | x |
|  | *Helminthotheca echioides* (L.) Holub | x |  |
|  | *Hypochaeris glabra* L. |  | x |
|  | *Hysterionica montevidensis* Baker |  | x |
|  | *Lucilia acutifolia* (Poir.) Cass | x |  |
|  | *Mikania cordifolia* (L.f.) Willd. |  | x |
|  | *Porophyllum ruderale* (Jacq.) Cass. |  | x |
|  | *Pseudognaphalium gaudichaudianum* (DC.) Anderb. |  | x |
|  | *Pterocaulon lorentzii* Malme | x | x |
|  | *Pterocaulon polystachyum* DC |  | x |
|  | *Senecio crassiflorus* (Poir.) DC. |  | x |
|  | *Senecio madagascariensis* Poir. |  | x |
|  | *Senecio selloi* (Spreng.) DC. |  | x |
|  | *Silybum marianum* (L.) Gaertn. |  | x |
|  | *Solidago chilensis* Meyen |  | x |
|  | *Sonchus asper* (L.) Hill |  | x |
|  | *Sonchus oleraceus* (L.) L. |  | x |
|  | *Tagetes minuta* L. |  | x |
|  | *Trixis pallida* Less. | x |  |
|  | *Vernonia incana* Less. | x | x |
|  | *Xanthium spinosum* L. |  | x |
| Convolvulaceae | *Dichondra macrocalyx* Meisn. |  | x |
|  | *Dichondra sericea* Sw*.* | x | x |
|  | *Evolvulus sericeus f. pedunculatus* Ooststr. | x | x |
| Cucurbitaceae | *Cayaponia sp.* |  | x |
| Cyperaceae | *Carex trachycystis* Griseb. |  | x |
|  | *Carex sp.* |  | x |
|  | *Cyperus sp.* | x | x |
|  | *Cyperus sp. 1* | x | x |
|  | *Cyperus sp. 2* |  | x |
|  | *Cyperus sp. 3* |  | x |
|  | *Cyperus sp. 4* |  | x |
|  | *Cyperus sp. 5* |  | x |
|  | *Fimbristylis autumnalis (L.)* Roem. & Schult. | x | x |
|  | *Kyllinga odorata* Vahl |  | x |
| Euphorbiaceae | *Acalypha communis* Müll.Arg. | x |  |
|  | *Astraea lobata* (L.) Klotzsch |  | x |
|  | *Bernardia sellowii* Müll.Arg. | x |  |
|  | *Croton sp.* | x |  |
|  | *Croton glandulosus L.* |  | x |
|  | *Euphorbia sp.* | x |  |
|  | *Euphorbia serpens* Kunth |  | x |
|  | *Tragia geraniifolia* Klotzsch ex Müll.Arg | x | x |
|  | *Tragia volubilis L.* |  | x |
| Gentianaceae | *Centaurium pulchellum* (Sw.) Druce |  | x |
| Geraniaceae | *Geranium carolinianum L.* |  | x |
| Hypericaceae | *Hypericum campestre* Cham. & Schltdl. |  | x |
| Iridaceae | *Sisyrinchium chilense Hook.* |  | x |
|  | *Sysirinchium sp.* |  | x |
| Juncaceae | *Juncus sp.* |  | x |
|  | *Juncus tenuis* Willd. |  | x |
| Lamiaceae | *Hyptis mutabilis* (Rich.) Briq. | x | x |
|  | *Teucrium vesicarium* Mill. |  | x |
| Leguminosae | *Acacia caven* (Molina) Molina |  | x |
|  | *Aeschynomene histrix var. incana* (Vogel) Benth. | x |  |
|  | *Astragalus cf. distinens Macloskie* |  | x |
|  | *Chamaecrista repens* (Vogel) H.S.Irwin & Barneby | x |  |
|  | *Desmodium affine* Schltdl. |  | x |
|  | *Desmodium sp.* |  | x |
|  | *Desmanthus virgatus* (L.) Willd. |  | x |
|  | *Lotus corniculatus* L. |  | x |
|  | *Lupinus multiflorus* Desr. | x |  |
|  | *Macroptilium prostratum* (Benth.) Urb*.* |  | x |
|  | *Prosopis affinis* Spreng |  | x |
|  | *Rhynchosia senna* Hook*.* |  | x |
|  | *Senna scabriuscula* (Vogel) H.S.Irwin & Barneby | x |  |
|  | *Stylosanthes sp.* | x |  |
|  | *Tephrosia cinerea* (L.) Pers. | x |  |
|  | *Zornia trachycarpa* Vogel | x |  |
| Lythraceae | *Cuphea glutinosa* Cham. & Schltdl. |  | x |
|  | *Heimia salicifolia* (Kunth) Link |  | x |
|  | *Lythrum maritimum* Kunth |  | x |
| Malvaceae | *Ayenia sp.* | x | x |
|  | *Krapovickasia flavescens* (Cav.) Fryxell |  | x |
|  | *Modiolastrum malvifolium* (Griseb.) K. Schum. |  | x |
|  | *Pavonia glechomoides* A. St.-Hil*.* |  | x |
|  | *Sida cf vespertina* Ekman |  | x |
|  | *Sida ciliaris L.* |  | x |
|  | *Sida rhombifolia L.* |  | x |
|  | *Sida spinosa L.* |  | x |
|  | *Wissadula glechomifolia* Hassl. |  | x |
| Meliaceae | *Melia azedarach L.* |  | x |
| Molluginaceae | *Mollugo verticillata L.* |  | x |
| Moraceae | *Dorstenia brasiliensis* Lam. | x |  |
| Myrtaceae | *Blepharocalyx salicifolius* (Kunth) O.Berg |  | x |
|  | *Eugenia myrcianthes* Nied. |  | x |
| Onagraceae | *Oenothera indecora* Cambess | x | x |
|  | *Oenothera parodiana* Munz | x |  |
|  | *Oenothera sp.1* |  | x |
| Orchidaceae | *Brachystele sp.* |  | x |
| Orobanchaceae | *Agalinis genistifolia* (Cham. & Schltdl.) D'Arcy |  | x |
| Oxalidaceae | *Oxalis conorrihiza* Jacq. |  | x |
|  | *Oxalis sellowiana* Zucc. | x |  |
|  | *Oxalis sp.* | x | x |
| Passifloraceae | *Passiflora sp.* | x |  |
| Plantaginaceae | *Kickxia elatine* (L.) Dumort*.* |  | x |
|  | *Plantago australis Lam.* | x | x |
|  | *Plantago brasiliensis* Sims | x |  |
|  | *Plantago tomentosa* Lam. |  | x |
|  | *Stemodia verticillata* (Mill.) Hassl. |  | x |
|  | *Veronica persica Poir.* |  | x |
| Poaceae | *Andropogon lateralis* Nees | x |  |
|  | *Andropogon sp.* | x |  |
|  | *Aristida circinalis* Lindm. | x |  |
|  | *Aristida sp.* | x |  |
|  | *Axonopus sp.* | x |  |
|  | *Bothriochloa laguroides* (DC.) Herter | x |  |
|  | *Briza sp.* | x | x |
|  | *Briza sp.1* | x | x |
|  | *Briza sp.2* |  | x |
|  | *Briza sp.3* |  | x |
|  | *Bromus catharticus* Vahl | x |  |
|  | *Calamagrostis sp.* |  | x |
|  | *Chloris* *elata* Desv. |  | x |
|  | *Chloris sp.* |  | x |
|  | *Cynodon dactylon (L.) Pers.* | x | x |
|  | *Digitaria aequiglumis* (Hack. & Arechav.) Parodi |  | x |
|  | *Digitaria sacchariflora* (Raddi) Henrard |  | x |
|  | *Echinochloa colona* (L.) Link | x | x |
|  | *Echinochloa sp.* |  | x |
|  | *Eleusine indica* (L.) Gaertn. |  | x |
|  | *Elionurus muticus* (Spreng.) Kuntze | x |  |
|  | *Eragrostis sp* |  | x |
|  | *Eragrostis sp1* |  | x |
|  | *Eragrostis sp2* |  | x |
|  | *Eustachys paspaloides* (Vahl) Lanza & Mattei | x | x |
|  | *Lolium sp.* | x |  |
|  | *Melica brasiliana Ard.* | x |  |
|  | *Panicum sabulorum Lam.* | x | x |
|  | *Panicum sp.* | x | x |
|  | *Paspalum* *dilatatum* Poir. |  | x |
|  | *Paspalum* *plicatulum* Michx. | x |  |
|  | *Paspalum distichum L.* |  | x |
|  | *Paspalum sp.* |  | x |
|  | *Paspalum sp.1* |  | x |
|  | *Piptochaetium montevidense* (Spreng.) *Parodi* | x | x |
|  | *Piptochaetium ruprechtianum É.Desv.* |  | x |
|  | *Piptochaetium sp.* |  | x |
|  | *Piptochaetium stipoides* (Trin. & Rupr.) Hack. & Arechav. |  | x |
|  | *Rottboellia sp.* | x |  |
|  | *Schizachyrium sp.* | x | x |
|  | *Setaria parviflora* (Poir.) M.Kerguelen | x | x |
|  | *Setaria sp.* |  | x |
|  | *Sporobolus sp.* | x |  |
|  | *Steinchisma hians* (Elliott) Nash |  | x |
|  | *Stipa sp.* | x |  |
|  | *Stipa sp.1* | x | x |
|  | *Stipa sp.2* | x | x |
| Polygalaceae | *Monnina* *cuneata* A. St.-Hil. | x | x |
|  | *Polygala duarteana A. St.-Hil. & Moq.* | x | x |
| Portulacaceae | *Portulaca grandiflora Hook.* |  | x |
|  | *Portulaca sp.* |  | x |
| Primulaceae | *Anagallis arvensis L.* |  | x |
| Ranunculaceae | *Clematis campestris A.St.-Hil.* |  | x |
| Rosaceae | *Margyricarpus pinnatus* (Lam.) Kuntze | x |  |
| Rubiaceae | *Galium aparine* L. | x |  |
|  | *Galium richardianum* (Gillies ex Hook. & Arn.) Endl. ex Walp. |  | x |
|  | *Mitracarpus megapotamicus* (Spreng.) *Kuntze* | x |  |
|  | *Richardia brasiliensis Gomes* | x | x |
|  | *Richardia humistrata* (Cham. & Schltdl.) *Steud.* | x | x |
|  | *Richardia stellaris* (Cham. & Schltdl.) Steud. |  | x |
|  | *Spermacoce glabra Michx.* |  | x |
|  | *Spermacoce verticillata L.* | x | x |
| Smilacaceae | *Smilax campestris Griseb.* |  | x |
| Solanaceae | *Bouchetia anomala* (Miers) Britton & Rusby |  | x |
|  | *Nicotiana longiflora Cav.* |  | x |
|  | *Petunia integrifolia* (Hook.) Schinz & Thell*.* | x | x |
|  | *Physalis angulata* L. |  | x |
|  | *Solanum atropurpureum* Schrank |  | x |
|  | *Solanum granuloso-leprosum* Dunal |  | x |
|  | *Solanum reflexum* Schrank |  | x |
|  | *Solanum sisymbriifolium Lam* |  | x |
|  | *Solanum sp.* |  | x |
|  | *Solanum sp1* |  | x |
| Urticaceae | *Parietaria debilis* G.Forst. |  | x |
| Verbenaceae | *Glandularia subincana* Tronc*.* |  | x |
|  | *Lantana megapotamica* (Spreng.) Tronc. | x |  |
|  | *Lippia cf arechavaletae* Moldenke | x |  |
|  | *Verbena bonariensis L.* |  | x |
|  | *Verbena gracilescens* (Cham.) Herter |  | x |
|  | *Verbena rigida* Spreng. |  | x |
| Violaceae | *Hybanthus parviflorus* (L.f.) Baill. | x | x |
