## appendixB for "Eucalyptus plantations in temperate grasslands: responses of taxonomic and functional diversity of plant communities to environmental changes"

**Appendix B.** Plant richness models selection results. Model rankings were based on AICc values. Variables are: CANOPY= Canopy cover, N= nitrogen content in soil, P= phosphorus content in soil, pH= soil pH, LITTER= leaf litter depth, TEMP= temperature at 10 cm above the ground.

| **CANOPY** | **N** | **P** | **pH** | **LITTER** | **TEMP** | **df** | **AICc** | **ΔAICc** | **weight** |
| --- | --- | --- | --- | --- | --- | --- | --- | --- | --- |
| - | - | - | -0.4767 | -0.1504 | - | 4 | 207.1667 | 0 | 0.2537 |
| - | 2.6849 | - | -0.5881 | -0.1532 | - | 5 | 208.0711 | 0.9044 | 0.1614 |
| - | - | - | - | -0.1198 | - | 3 | 209.0708 | 1.9041 | 0.0979 |
| - | - | - | -0.4687 | -0.1529 | -0.0160 | 5 | 210.1617 | 2.9950 | 0.0567 |
| 0.0004 | - | - | -0.4719 | -0.1514 | - | 5 | 210.1975 | 3.0307 | 0.0557 |
| - | - | -0.0034 | -0.4712 | -0.1502 | - | 5 | 210.1993 | 3.0326 | 0.0557 |
| - | 2.8275 | - | -0.5741 | -0.1586 | -0.0353 | 6 | 211.1838 | 4.0170 | 0.0340 |
| - | - | -0.0334 | - | -0.1203 | - | 4 | 211.3045 | 4.1378 | 0.0320 |
| - | 2.7007 | -0.0062 | -0.5797 | -0.1528 | - | 6 | 211.3908 | 4.2241 | 0.0307 |
| 0.0001 | 2.6823 | - | -0.5861 | -0.1536 | - | 6 | 211.4128 | 4.2461 | 0.0304 |
| 0.0021 | - | - | - | -0.1266 | - | 4 | 211.6252 | 4.4585 | 0.0273 |
| - | 0.9359 | - | - | -0.1180 | - | 4 | 211.6295 | 4.4628 | 0.0272 |
| - | - | - | - | -0.1259 | -0.0340 | 4 | 211.6739 | 4.5072 | 0.0266 |
| -0.0005 | - | - | -0.4717 | -0.1526 | -0.0225 | 6 | 213.4976 | 6.3309 | 0.0107 |
| - | - | -0.0032 | -0.4635 | -0.1527 | -0.0159 | 6 | 213.4990 | 6.3323 | 0.0107 |
| 0.0004 | - | -0.0036 | -0.4659 | -0.1512 | - | 6 | 213.5333 | 6.3666 | 0.0105 |
| - | 1.0920 | -0.0359 | - | -0.1181 | - | 5 | 214.0431 | 6.8764 | 0.0081 |
| 0.0022 | - | -0.0336 | - | -0.1274 | - | 5 | 214.1075 | 6.9408 | 0.0079 |
| - | - | -0.0333 | - | -0.1265 | -0.0336 | 5 | 214.1716 | 7.0049 | 0.0076 |
| - | 1.1710 | - | - | -0.1254 | -0.0443 | 5 | 214.3857 | 7.2190 | 0.0069 |
| 0.0023 | 1.0095 | - | - | -0.1251 | - | 5 | 214.4113 | 7.2446 | 0.0068 |
| 0.0016 | - | - | - | -0.1272 | -0.0128 | 5 | 214.6525 | 7.4858 | 0.0060 |
| -0.0027 | 3.0150 | - | -0.5988 | -0.1573 | -0.0707 | 7 | 214.6705 | 7.5038 | 0.0060 |
| - | 2.8457 | -0.0064 | -0.5653 | -0.1582 | -0.0354 | 7 | 214.8539 | 7.6872 | 0.0054 |
| 0.0002 | 2.6977 | -0.0063 | -0.5769 | -0.1532 | - | 7 | 215.0836 | 7.9169 | 0.0048 |
| - | - | - | - | - | - | 2 | 216.5491 | 9.3824 | 0.0023 |
| - | 1.3652 | -0.0368 | - | -0.1260 | -0.0467 | 6 | 217.0656 | 9.8989 | 0.0018 |
| 0.0025 | 1.1948 | -0.0367 | - | -0.1258 | - | 6 | 217.0905 | 9.9238 | 0.0018 |
| -0.0005 | - | -0.0030 | -0.4668 | -0.1524 | -0.0219 | 7 | 217.1877 | 10.0210 | 0.0017 |
| 0.0018 | - | -0.0335 | - | -0.1280 | -0.0103 | 6 | 217.4428 | 10.2761 | 0.0015 |
| 0.0012 | 1.1265 | - | - | -0.1264 | -0.0287 | 6 | 217.6975 | 10.5307 | 0.0013 |
| - | - | - | - | - | 0.0724 | 3 | 218.3821 | 11.2154 | 0.0009 |
| -0.0036 | - | - | - | - | - | 3 | 218.5210 | 11.3543 | 0.0009 |
| - | 1.7264 | - | - | - | - | 3 | 218.5954 | 11.4287 | 0.0008 |
| - | - | -0.0313 | - | - | - | 3 | 218.7567 | 11.5900 | 0.0008 |
| -0.0027 | 3.0251 | -0.0050 | -0.5915 | -0.1570 | -0.0702 | 8 | 218.7608 | 11.5941 | 0.0008 |
| - | - | - | -0.1245 | - | - | 3 | 218.8400 | 11.6733 | 0.0007 |
| - | - | - | -0.1869 | - | 0.0865 | 4 | 220.6473 | 13.4806 | 0.0003 |
| - | 2.4978 | - | -0.2200 | - | - | 4 | 220.6707 | 13.5040 | 0.0003 |
| -0.0047 | - | - | -0.2061 | - | - | 4 | 220.6955 | 13.5288 | 0.0003 |
| 0.0013 | 1.3198 | -0.0368 | - | -0.1271 | -0.0298 | 7 | 220.7243 | 13.5576 | 0.0003 |
| - | - | -0.0330 | - | - | 0.0738 | 4 | 220.7759 | 13.6092 | 0.0003 |
| - | 1.9609 | -0.0377 | - | - | - | 4 | 220.8838 | 13.7171 | 0.0003 |
| -0.0032 | 1.5237 | - | - | - | - | 4 | 220.9047 | 13.7380 | 0.0003 |
| - | 1.2208 | - | - | - | 0.0595 | 4 | 220.9265 | 13.7598 | 0.0003 |
| -0.0036 | - | -0.0324 | - | - | - | 4 | 220.9321 | 13.7654 | 0.0003 |
| -0.0014 | - | - | - | - | 0.0542 | 4 | 221.1197 | 13.9530 | 0.0002 |
| - | - | -0.0248 | -0.0847 | - | - | 4 | 221.4275 | 14.2608 | 0.0002 |
| -0.0049 | 2.5322 | - | -0.3141 | - | - | 5 | 222.7263 | 15.5595 | 0.0001 |
| - | 2.0151 | - | -0.2552 | - | 0.0704 | 5 | 223.0910 | 15.9243 | 0.0001 |
| - | 2.5721 | -0.0272 | -0.1842 | - | - | 5 | 223.4703 | 16.3036 | 0.0001 |
| -0.0032 | 1.7153 | -0.0369 | - | - | - | 5 | 223.4709 | 16.3042 | 0.0001 |
| - | 1.4254 | -0.0365 | - | - | 0.0577 | 5 | 223.5007 | 16.3340 | 0.0001 |
| - | - | -0.0233 | -0.1518 | - | 0.0857 | 5 | 223.5092 | 16.3425 | 0.0001 |
| -0.0026 | - | - | -0.2084 | - | 0.0538 | 5 | 223.5571 | 16.3904 | 0.0001 |
| -0.0046 | - | -0.0209 | -0.1722 | - | - | 5 | 223.5956 | 16.4289 | 0.0001 |
| -0.0013 | - | -0.0328 | - | - | 0.0563 | 5 | 223.7818 | 16.6151 | 0.0001 |
| -0.0020 | 1.3366 | - | - | - | 0.0318 | 5 | 223.8880 | 16.7213 | 0.0001 |
| -0.0047 | 2.5678 | -0.0220 | -0.2826 | - | - | 6 | 225.9057 | 18.7390 | 0.0000 |
| -0.0045 | 2.4608 | - | -0.3115 | - | 0.0105 | 6 | 226.0629 | 18.8962 | 0.0000 |
| - | 2.0747 | -0.0248 | -0.2225 | - | 0.0685 | 6 | 226.2265 | 19.0598 | 0.0000 |
| -0.0023 | - | -0.0217 | -0.1737 | - | 0.0560 | 6 | 226.7477 | 19.5810 | 0.0000 |
| -0.0020 | 1.5361 | -0.0366 | - | - | 0.0295 | 6 | 226.7661 | 19.5994 | 0.0000 |
| -0.0043 | 2.4879 | -0.0221 | -0.2796 | - | 0.0116 | 7 | 229.5929 | 22.4262 | 0.0000 |
