## appendixC for "Eucalyptus plantations in temperate grasslands: responses of taxonomic and functional diversity of plant communities to environmental changes"

**Appendix C.** Taxonomic similarity models selection results. Model rankings were based on AICc values. Variables are: sCANOPY= similarity in canopy cover, sN= similarity in nitrogen content in soil, sP= similarity in phosphorus content in soil, spH= similarity in soil pH, sLITTER= similarity in leaf litter depth, sTEMP= similarity in temperature at 10 cm above the ground.

| **sCANOPY** | **sLITTER** | **sN** | **sP** | **spH** | **sTEMP** | **df** | **AICc** | **ΔAICc** | **weight** |
| --- | --- | --- | --- | --- | --- | --- | --- | --- | --- |
| - | 0.748 | - | - | - | - | 3 | -68.272 | 0.000 | 0.174 |
| -0.474 | 1.017 | - | - | - | - | 4 | -67.309 | 0.963 | 0.108 |
| - | 0.760 | -0.302 | - | - | - | 4 | -66.165 | 2.107 | 0.061 |
| -0.936 | 0.960 | - | - | -0.994 | - | 5 | -65.933 | 2.338 | 0.054 |
| - | 0.863 | - | - | - | -0.201 | 4 | -65.852 | 2.420 | 0.052 |
| - | 0.731 | - | -0.452 | - | - | 4 | -65.702 | 2.570 | 0.048 |
| - | - | - | - | - | - | 2 | -65.665 | 2.606 | 0.047 |
| - | 0.764 | - | - | 0.046 | - | 4 | -65.504 | 2.768 | 0.044 |
| -0.459 | 1.022 | -0.284 | - | - | - | 5 | -64.897 | 3.375 | 0.032 |
| -0.721 | 0.967 | - | - | - | 0.334 | 5 | -64.679 | 3.593 | 0.029 |
| -0.496 | 1.002 | - | -0.589 | - | - | 5 | -64.645 | 3.626 | 0.028 |
| - | - | - | - | -0.547 | - | 3 | -64.132 | 4.139 | 0.022 |
| - | 0.930 | -0.374 | - | - | -0.282 | 5 | -63.774 | 4.498 | 0.018 |
| - | - | -0.279 | - | - | - | 3 | -63.588 | 4.684 | 0.017 |
| - | - | - | -0.851 | - | - | 3 | -63.562 | 4.710 | 0.017 |
| - | - | - | - | - | 0.223 | 3 | -63.519 | 4.753 | 0.016 |
| - | 0.751 | -0.322 | -0.532 | - | - | 5 | -63.416 | 4.856 | 0.015 |
| - | 0.840 | - | - | -0.523 | -0.460 | 5 | -63.174 | 5.098 | 0.014 |
| 0.056 | - | - | - | - | - | 3 | -63.143 | 5.128 | 0.013 |
| - | 0.777 | -0.303 | - | 0.048 | - | 5 | -63.134 | 5.138 | 0.013 |
| -0.892 | 0.955 | -0.240 | - | -0.954 | - | 6 | -63.095 | 5.176 | 0.013 |
| - | 0.849 | - | -0.483 | - | -0.211 | 5 | -63.056 | 5.215 | 0.013 |
| -0.924 | 0.952 | - | -0.515 | -0.939 | - | 6 | -62.863 | 5.408 | 0.012 |
| - | 0.752 | - | -0.459 | 0.063 | - | 5 | -62.677 | 5.595 | 0.011 |
| -0.918 | 0.970 | - | - | -1.047 | -0.062 | 6 | -62.600 | 5.672 | 0.010 |
| -0.530 | - | - | - | -1.171 | - | 4 | -62.245 | 6.026 | 0.009 |
| -0.469 | 1.011 | -0.283 | -0.580 | - | - | 6 | -61.929 | 6.343 | 0.007 |
| - | - | -0.297 | - | -0.574 | - | 4 | -61.929 | 6.343 | 0.007 |
| -0.615 | 0.986 | -0.227 | - | - | 0.210 | 6 | -61.691 | 6.581 | 0.006 |
| -0.723 | 0.953 | - | -0.579 | - | 0.316 | 6 | -61.685 | 6.586 | 0.006 |
| - | - | - | -0.640 | -0.500 | - | 4 | -61.615 | 6.656 | 0.006 |
| - | - | - | - | -0.712 | -0.130 | 4 | -61.413 | 6.859 | 0.006 |
| -0.540 | - | - | - | - | 0.676 | 4 | -61.309 | 6.962 | 0.005 |
| - | - | -0.262 | -0.796 | - | - | 4 | -61.202 | 7.069 | 0.005 |
| - | 0.916 | -0.434 | - | -0.732 | -0.653 | 6 | -61.145 | 7.126 | 0.005 |
| - | - | -0.254 | - | - | 0.202 | 4 | -61.136 | 7.135 | 0.005 |
| - | - | - | -0.753 | - | 0.196 | 4 | -61.095 | 7.177 | 0.005 |
| 0.066 | - | -0.282 | - | - | - | 4 | -60.843 | 7.428 | 0.004 |
| 0.011 | - | - | -0.843 | - | - | 4 | -60.788 | 7.484 | 0.004 |
| - | 0.922 | -0.385 | -0.536 | - | -0.285 | 6 | -60.747 | 7.524 | 0.004 |
| - | 0.771 | -0.322 | -0.534 | 0.058 | - | 6 | -60.085 | 8.187 | 0.003 |
| - | 0.831 | - | -0.444 | -0.482 | -0.448 | 6 | -60.034 | 8.237 | 0.003 |
| -0.512 | - | -0.284 | - | -1.180 | - | 5 | -59.767 | 8.504 | 0.002 |
| -0.870 | 0.952 | -0.238 | -0.498 | -0.897 | - | 7 | -59.664 | 8.608 | 0.002 |
| -0.804 | 0.997 | -0.284 | - | -1.144 | -0.243 | 7 | -59.532 | 8.739 | 0.002 |
| -0.519 | - | - | -0.672 | -1.098 | - | 5 | -59.472 | 8.800 | 0.002 |
| -0.756 | - | - | - | -0.983 | 0.364 | 5 | -59.445 | 8.826 | 0.002 |
| -0.908 | 0.962 | - | -0.513 | -0.988 | -0.056 | 7 | -59.176 | 9.095 | 0.002 |
| - | - | -0.297 | -0.620 | -0.543 | - | 5 | -59.145 | 9.127 | 0.002 |
| - | - | -0.325 | - | -0.842 | -0.211 | 5 | -59.040 | 9.231 | 0.002 |
| -0.557 | - | - | -0.867 | - | 0.654 | 5 | -58.707 | 9.565 | 0.001 |
| - | - | - | -0.629 | -0.641 | -0.113 | 5 | -58.620 | 9.651 | 0.001 |
| -0.457 | - | -0.190 | - | - | 0.592 | 5 | -58.486 | 9.786 | 0.001 |
| - | - | -0.252 | -0.744 | - | 0.188 | 5 | -58.450 | 9.822 | 0.001 |
| -0.613 | 0.976 | -0.231 | -0.579 | - | 0.196 | 7 | -58.360 | 9.911 | 0.001 |
| 0.041 | - | -0.267 | -0.777 | - | - | 5 | -58.175 | 10.097 | 0.001 |
| - | 0.913 | -0.437 | -0.471 | -0.676 | -0.623 | 7 | -57.694 | 10.578 | 0.001 |
| -0.486 | - | -0.269 | -0.564 | -1.110 | - | 6 | -56.620 | 11.651 | 0.001 |
| -0.668 | - | -0.253 | - | -1.062 | 0.241 | 6 | -56.528 | 11.744 | 0.000 |
| -0.749 | - | - | -0.713 | -0.901 | 0.375 | 6 | -56.399 | 11.873 | 0.000 |
| - | - | -0.315 | -0.574 | -0.764 | -0.175 | 6 | -55.910 | 12.361 | 0.000 |
| -0.791 | 0.993 | -0.279 | -0.479 | -1.071 | -0.224 | 8 | -55.675 | 12.596 | 0.000 |
| -0.474 | - | -0.177 | -0.825 | - | 0.583 | 6 | -55.551 | 12.720 | 0.000 |
| -0.659 | - | -0.234 | -0.615 | -0.973 | 0.270 | 7 | -53.062 | 15.210 | 0.000 |
