## appendixD for "Eucalyptus plantations in temperate grasslands: responses of taxonomic and functional diversity of plant communities to environmental changes"

**Appendix D.** Alpha functional diversity models selection results. Model rankings were based on AICc values. Variables are: CANOPY= Canopy cover, C= carbon content in soil, N= nitrogen content in soil, P= phosphorus content in soil, pH= soil pH, LITTER= leaf litter depth, TEMP= temperature at 10 cm above the ground.

| **CANOPY** | **C** | **LITTER** | **N** | **P** | **pH** | **TEMP** | **df** | **AICc** | **ΔAICc** | **weight** |
| --- | --- | --- | --- | --- | --- | --- | --- | --- | --- | --- |
| -0.0010 | - | - | - | - | - | - | 3 | -87.3767 | 0 | 0.1628 |
| -0.0008 | - | -0.0034 | - | - | - | - | 4 | -85.9271 | 1.4496 | 0.0789 |
| - | - | -0.0063 | - | - | - | - | 3 | -84.8373 | 2.5394 | 0.0457 |
| -0.0010 | - | - | - | 0.0012 | - | - | 4 | -84.7364 | 2.6403 | 0.0435 |
| -0.0011 | - | - | - | - | - | -0.0027 | 4 | -84.6799 | 2.6968 | 0.0423 |
| -0.0010 | - | - | 0.0303 | - | - | - | 4 | -84.6447 | 2.7319 | 0.0415 |
| -0.0010 | -0.0022 | - | - | - | - | - | 4 | -84.6368 | 2.7399 | 0.0414 |
| -0.0009 | - | - | - | - | 0.0009 | - | 4 | -84.6047 | 2.7720 | 0.0407 |
| - | - | - | - | - | - | 0.0126 | 3 | -83.8400 | 3.5367 | 0.0278 |
| - | - | -0.0047 | - | - | - | 0.0078 | 4 | -83.3126 | 4.0640 | 0.0213 |
| -0.0007 | -0.0060 | -0.0038 | - | - | - | - | 5 | -83.1367 | 4.2400 | 0.0195 |
| -0.0009 | - | -0.0037 | - | - | - | -0.0045 | 5 | -83.1075 | 4.2691 | 0.0193 |
| -0.0008 | - | -0.0034 | - | 0.0012 | - | - | 5 | -83.0275 | 4.3492 | 0.0185 |
| -0.0008 | - | -0.0038 | - | - | -0.0058 | - | 5 | -82.9913 | 4.3854 | 0.0182 |
| -0.0008 | - | -0.0034 | 0.0202 | - | - | - | 5 | -82.9078 | 4.4688 | 0.0174 |
| - | - | - | - | - | - | - | 2 | -82.7552 | 4.6215 | 0.0162 |
| - | -0.0090 | -0.0067 | - | - | - | - | 4 | -82.5627 | 4.8140 | 0.0147 |
| - | - | -0.0058 | - | - | 0.0066 | - | 4 | -82.1935 | 5.1832 | 0.0122 |
| - | - | -0.0063 | - | 0.0011 | - | - | 4 | -82.1564 | 5.2202 | 0.0120 |
| - | - | -0.0063 | 0.0103 | - | - | - | 4 | -82.0671 | 5.3096 | 0.0114 |
| - | - | - | - | - | 0.0221 | - | 3 | -81.8329 | 5.5437 | 0.0102 |
| -0.0011 | - | - | - | 0.0013 | - | -0.0028 | 5 | -81.7810 | 5.5957 | 0.0099 |
| -0.0010 | -0.0064 | - | 0.0812 | - | - | - | 5 | -81.7790 | 5.5977 | 0.0099 |
| -0.0010 | -0.0025 | - | - | 0.0013 | - | - | 5 | -81.7429 | 5.6338 | 0.0097 |
| -0.0010 | - | - | 0.0239 | 0.0012 | - | - | 5 | -81.7238 | 5.6529 | 0.0096 |
| -0.0011 | - | - | 0.0401 | - | - | -0.0033 | 5 | -81.7129 | 5.6638 | 0.0096 |
| -0.0010 | - | - | - | 0.0013 | -0.0007 | - | 5 | -81.6988 | 5.6779 | 0.0095 |
| -0.0011 | - | - | - | - | 0.0015 | -0.0028 | 5 | -81.6483 | 5.7283 | 0.0093 |
| -0.0011 | -0.0010 | - | - | - | - | -0.0024 | 5 | -81.6467 | 5.7300 | 0.0093 |
| -0.0010 | - | - | 0.0305 | - | -0.0001 | - | 5 | -81.6058 | 5.7709 | 0.0091 |
| -0.0009 | -0.0023 | - | - | - | 0.0012 | - | 5 | -81.6026 | 5.7741 | 0.0091 |
| - | -0.0094 | - | - | - | - | 0.0137 | 4 | -81.5794 | 5.7973 | 0.0090 |
| - | - | - | - | - | 0.0095 | 0.0109 | 4 | -81.3320 | 6.0447 | 0.0079 |
| - | -0.0118 | -0.0051 | - | - | - | 0.0089 | 5 | -81.1551 | 6.2216 | 0.0073 |
| - | - | - | - | 0.0010 | - | 0.0126 | 4 | -81.1480 | 6.2287 | 0.0072 |
| - | - | - | -0.0082 | - | - | 0.0126 | 4 | -81.0679 | 6.3087 | 0.0069 |
| - | - | -0.0047 | - | 0.0011 | - | 0.0078 | 5 | -80.3716 | 7.0051 | 0.0049 |
| - | - | - | - | 0.0010 | - | - | 3 | -80.2804 | 7.0963 | 0.0047 |
| - | - | -0.0047 | -0.0077 | - | - | 0.0078 | 5 | -80.2763 | 7.1004 | 0.0047 |
| - | - | -0.0047 | - | - | 0.0006 | 0.0077 | 5 | -80.2749 | 7.1018 | 0.0047 |
| - | -0.0025 | - | - | - | - | - | 3 | -80.2467 | 7.1300 | 0.0046 |
| - | - | - | 0.0281 | - | - | - | 3 | -80.2398 | 7.1368 | 0.0046 |
| -0.0007 | -0.0121 | -0.0041 | 0.1140 | - | - | - | 6 | -80.1669 | 7.2098 | 0.0044 |
| - | -0.0164 | -0.0070 | 0.1375 | - | - | - | 5 | -80.0016 | 7.3751 | 0.0041 |
| -0.0007 | -0.0063 | -0.0038 | - | 0.0014 | - | - | 6 | -79.9635 | 7.4132 | 0.0040 |
| -0.0009 | - | -0.0037 | - | 0.0013 | - | -0.0046 | 6 | -79.9141 | 7.4626 | 0.0039 |
| -0.0008 | -0.0059 | -0.0041 | - | - | -0.0057 | - | 6 | -79.8944 | 7.4823 | 0.0039 |
| -0.0009 | -0.0044 | -0.0039 | - | - | - | -0.0031 | 6 | -79.8802 | 7.4965 | 0.0038 |
| -0.0008 | - | -0.0039 | - | 0.0016 | -0.0080 | - | 6 | -79.8695 | 7.5072 | 0.0038 |
| -0.0010 | - | -0.0039 | - | - | -0.0052 | -0.0043 | 6 | -79.8475 | 7.5292 | 0.0038 |
| -0.0010 | - | -0.0036 | 0.0344 | - | - | -0.0050 | 6 | -79.8204 | 7.5563 | 0.0037 |
| -0.0008 | - | -0.0038 | 0.0346 | - | -0.0070 | - | 6 | -79.7027 | 7.6740 | 0.0035 |
| -0.0008 | - | -0.0034 | 0.0136 | 0.0012 | - | - | 6 | -79.6936 | 7.6831 | 0.0035 |
| - | -0.0093 | -0.0068 | - | 0.0013 | - | - | 5 | -79.6551 | 7.7215 | 0.0034 |
| - | -0.0089 | -0.0062 | - | - | 0.0062 | - | 5 | -79.6405 | 7.7362 | 0.0034 |
| - | - | -0.0058 | - | 0.0008 | 0.0056 | - | 5 | -79.2075 | 8.1691 | 0.0027 |
| - | - | -0.0058 | -0.0029 | - | 0.0067 | - | 5 | -79.1549 | 8.2218 | 0.0027 |
| - | -0.0038 | - | - | - | 0.0225 | - | 4 | -79.1401 | 8.2366 | 0.0026 |
| - | - | -0.0063 | 0.0044 | 0.0011 | - | - | 5 | -79.1183 | 8.2584 | 0.0026 |
| - | - | - | -0.0217 | - | 0.0227 | - | 4 | -79.0750 | 8.3016 | 0.0026 |
| - | - | - | - | 0.0001 | 0.0221 | - | 4 | -79.0584 | 8.3182 | 0.0025 |
| - | -0.0150 | - | 0.1064 | - | - | 0.0139 | 5 | -78.8179 | 8.5587 | 0.0023 |
| - | -0.0091 | - | - | - | 0.0089 | 0.0122 | 5 | -78.7792 | 8.5975 | 0.0022 |
| - | -0.0097 | - | - | 0.0012 | - | 0.0138 | 5 | -78.6587 | 8.7179 | 0.0021 |
| -0.0010 | -0.0064 | - | 0.0743 | 0.0012 | - | - | 6 | -78.5506 | 8.8261 | 0.0020 |
| -0.0011 | - | - | 0.0337 | 0.0012 | - | -0.0033 | 6 | -78.4883 | 8.8884 | 0.0019 |
| -0.0010 | -0.0053 | - | 0.0777 | - | - | -0.0019 | 6 | -78.4683 | 8.9084 | 0.0019 |
| -0.0011 | -0.0013 | - | - | 0.0013 | - | -0.0024 | 6 | -78.4483 | 8.9284 | 0.0019 |
| -0.0010 | -0.0065 | - | 0.0839 | - | -0.0009 | - | 6 | -78.4387 | 8.9380 | 0.0019 |
| -0.0011 | - | - | - | 0.0013 | -0.0001 | -0.0028 | 6 | -78.4381 | 8.9386 | 0.0019 |
| -0.0010 | -0.0025 | - | - | 0.0013 | -0.0004 | - | 6 | -78.4005 | 8.9762 | 0.0018 |
| -0.0010 | - | - | 0.0269 | 0.0012 | -0.0015 | - | 6 | -78.3871 | 8.9896 | 0.0018 |
| -0.0011 | - | - | 0.0394 | - | 0.0004 | -0.0033 | 6 | -78.3704 | 9.0063 | 0.0018 |
| - | -0.0193 | -0.0053 | 0.1399 | - | - | 0.0090 | 6 | -78.3378 | 9.0389 | 0.0018 |
| - | - | - | - | 0.0007 | 0.0088 | 0.0111 | 5 | -78.3247 | 9.0520 | 0.0018 |
| - | - | - | -0.0256 | - | 0.0101 | 0.0109 | 5 | -78.3185 | 9.0582 | 0.0018 |
| -0.0010 | -0.0010 | - | - | - | 0.0016 | -0.0025 | 6 | -78.3118 | 9.0649 | 0.0018 |
| - | - | - | -0.0142 | 0.0011 | - | 0.0126 | 5 | -78.1170 | 9.2597 | 0.0016 |
| - | -0.0122 | -0.0051 | - | 0.0013 | - | 0.0089 | 6 | -77.9641 | 9.4125 | 0.0015 |
| - | -0.0119 | -0.0051 | - | - | -0.0009 | 0.0090 | 6 | -77.8148 | 9.5619 | 0.0014 |
| - | -0.0068 | - | 0.0817 | - | - | - | 4 | -77.6125 | 9.7642 | 0.0012 |
| - | -0.0028 | - | - | 0.0011 | - | - | 4 | -77.5482 | 9.8285 | 0.0012 |
| - | - | - | 0.0229 | 0.0009 | - | - | 4 | -77.5242 | 9.8525 | 0.0012 |
| - | - | -0.0047 | -0.0140 | 0.0011 | - | 0.0078 | 6 | -77.0371 | 10.3396 | 0.0009 |
| - | - | -0.0048 | - | 0.0011 | -0.0008 | 0.0079 | 6 | -77.0304 | 10.3463 | 0.0009 |
| - | - | -0.0047 | -0.0093 | - | 0.0009 | 0.0077 | 6 | -76.9356 | 10.4411 | 0.0009 |
| -0.0008 | -0.0140 | -0.0047 | 0.1505 | - | -0.0105 | - | 7 | -76.7820 | 10.5947 | 0.0008 |
| - | -0.0163 | -0.0070 | 0.1314 | 0.0010 | - | - | 6 | -76.7454 | 10.6313 | 0.0008 |
| - | -0.0159 | -0.0068 | 0.1292 | - | 0.0023 | - | 6 | -76.6728 | 10.7039 | 0.0008 |
| -0.0007 | -0.0121 | -0.0041 | 0.1071 | 0.0012 | - | - | 7 | -76.5958 | 10.7809 | 0.0007 |
| -0.0008 | -0.0107 | -0.0041 | 0.1097 | - | - | -0.0025 | 7 | -76.5289 | 10.8478 | 0.0007 |
| -0.0008 | -0.0064 | -0.0043 | - | 0.0017 | -0.0081 | - | 7 | -76.4587 | 10.9179 | 0.0007 |
| - | -0.0092 | -0.0063 | - | 0.0011 | 0.0049 | - | 6 | -76.3827 | 10.9940 | 0.0007 |
| -0.0010 | - | -0.0040 | - | 0.0016 | -0.0074 | -0.0043 | 7 | -76.3779 | 10.9988 | 0.0007 |
| -0.0009 | -0.0048 | -0.0039 | - | 0.0014 | - | -0.0031 | 7 | -76.3546 | 11.0221 | 0.0007 |
| -0.0009 | -0.0045 | -0.0042 | - | - | -0.0053 | -0.0029 | 7 | -76.2729 | 11.1038 | 0.0006 |
| -0.0010 | - | -0.0036 | 0.0278 | 0.0012 | - | -0.0050 | 7 | -76.2555 | 11.1212 | 0.0006 |
| -0.0010 | - | -0.0040 | 0.0480 | - | -0.0067 | -0.0049 | 7 | -76.2548 | 11.1219 | 0.0006 |
| -0.0008 | - | -0.0039 | 0.0302 | 0.0016 | -0.0089 | - | 7 | -76.2162 | 11.1604 | 0.0006 |
| - | -0.0046 | - | 0.0155 | - | 0.0222 | - | 5 | -76.1062 | 11.2705 | 0.0006 |
| - | -0.0038 | - | - | 0.0001 | 0.0224 | - | 5 | -76.1021 | 11.2746 | 0.0006 |
| - | - | - | -0.0220 | 0.0001 | 0.0226 | - | 5 | -76.0367 | 11.3400 | 0.0006 |
| - | - | -0.0058 | -0.0058 | 0.0008 | 0.0058 | - | 6 | -75.8660 | 11.5107 | 0.0005 |
| - | -0.0136 | - | 0.0844 | - | 0.0066 | 0.0127 | 6 | -75.5966 | 11.7801 | 0.0005 |
| - | -0.0149 | - | 0.1002 | 0.0010 | - | 0.0140 | 6 | -75.5600 | 11.8167 | 0.0004 |
| - | -0.0093 | - | - | 0.0009 | 0.0079 | 0.0124 | 6 | -75.4940 | 11.8827 | 0.0004 |
| - | - | - | -0.0282 | 0.0007 | 0.0094 | 0.0111 | 6 | -75.0127 | 12.3640 | 0.0003 |
| -0.0010 | -0.0052 | - | 0.0707 | 0.0012 | - | -0.0020 | 7 | -74.8888 | 12.4878 | 0.0003 |
| -0.0010 | -0.0066 | - | 0.0808 | 0.0012 | -0.0024 | - | 7 | -74.8719 | 12.5048 | 0.0003 |
| -0.0011 | - | - | 0.0357 | 0.0012 | -0.0010 | -0.0032 | 7 | -74.7967 | 12.5800 | 0.0003 |
| -0.0010 | -0.0053 | - | 0.0793 | - | -0.0005 | -0.0019 | 7 | -74.7744 | 12.6023 | 0.0003 |
| -0.0011 | -0.0013 | - | - | 0.0013 | 0.0000 | -0.0024 | 7 | -74.7535 | 12.6231 | 0.0003 |
| - | -0.0210 | -0.0057 | 0.1641 | - | -0.0065 | 0.0097 | 7 | -74.7533 | 12.6234 | 0.0003 |
| - | -0.0193 | -0.0053 | 0.1335 | 0.0011 | - | 0.0090 | 7 | -74.7447 | 12.6320 | 0.0003 |
| - | -0.0067 | - | 0.0761 | 0.0009 | - | - | 5 | -74.6301 | 12.7465 | 0.0003 |
| - | -0.0124 | -0.0052 | - | 0.0015 | -0.0029 | 0.0093 | 7 | -74.2926 | 13.0841 | 0.0002 |
| - | - | -0.0047 | -0.0133 | 0.0011 | -0.0005 | 0.0079 | 7 | -73.3430 | 14.0337 | 0.0001 |
| - | -0.0161 | -0.0069 | 0.1275 | 0.0010 | 0.0011 | - | 7 | -73.0540 | 14.3227 | 0.0001 |
| -0.0008 | -0.0143 | -0.0048 | 0.1481 | 0.0017 | -0.0128 | - | 8 | -72.9227 | 14.4540 | 0.0001 |
| - | -0.0046 | - | 0.0151 | 0.0001 | 0.0221 | - | 6 | -72.7640 | 14.6127 | 0.0001 |
| -0.0009 | -0.0129 | -0.0047 | 0.1460 | - | -0.0102 | -0.0018 | 8 | -72.7075 | 14.6692 | 0.0001 |
| -0.0008 | -0.0106 | -0.0041 | 0.1027 | 0.0012 | - | -0.0025 | 8 | -72.5485 | 14.8282 | 0.0001 |
| -0.0009 | -0.0050 | -0.0043 | - | 0.0017 | -0.0077 | -0.0027 | 8 | -72.4207 | 14.9560 | 0.0001 |
| -0.0010 | - | -0.0041 | 0.0435 | 0.0015 | -0.0087 | -0.0049 | 8 | -72.3569 | 15.0198 | 0.0001 |
| - | -0.0138 | - | 0.0825 | 0.0008 | 0.0057 | 0.0129 | 7 | -71.9523 | 15.4244 | 0.0001 |
| - | -0.0214 | -0.0058 | 0.1626 | 0.0014 | -0.0084 | 0.0100 | 8 | -70.8126 | 16.5641 | 0.0000 |
| -0.0011 | -0.0054 | - | 0.0764 | 0.0012 | -0.0020 | -0.0018 | 8 | -70.7945 | 16.5822 | 0.0000 |
| -0.0009 | -0.0132 | -0.0048 | 0.1439 | 0.0017 | -0.0124 | -0.0017 | 9 | -68.3618 | 19.0149 | 0.0000 |
