## appendixE for "Eucalyptus plantations in temperate grasslands: responses of taxonomic and functional diversity of plant communities to environmental changes"

**Appendix E.** Plant functional similarity models selection results. Model rankings were based on AICc values. Variables are: sCANOPY= similarity in canopy cover, sN= similarity in nitrogen content in soil, sP= similarity in phosphorus content in soil, spH= similarity in soil pH, sLITTER= similarity in leaf litter depth, sTEMP= similarity in temperature at 10 cm above the ground.

| **sCANOPY** | **sLITTER** | **sN** | **sP** | **spH** | **sTEMP** | **df** | **AICc** | **ΔAICc** | **weight** |
| --- | --- | --- | --- | --- | --- | --- | --- | --- | --- |
| - | 0.406 | - | - | - | - | 3 | -34.400 | 0.000 | 0.173 |
| - | 0.350 | 0.290 | - | -0.559 | - | 5 | -33.268 | 1.131 | 0.098 |
| - | 0.353 | - | - | -0.328 | - | 4 | -33.187 | 1.213 | 0.094 |
| - | 0.412 | 0.153 | - | - | - | 4 | -32.547 | 1.853 | 0.069 |
| - | 0.360 | - | - | - | 0.101 | 4 | -32.128 | 2.271 | 0.056 |
| - | 0.405 | - | 0.044 | - | - | 4 | -31.601 | 2.799 | 0.043 |
| -0.001 | 0.407 | - | - | - | - | 4 | -31.586 | 2.814 | 0.042 |
| -0.184 | 0.413 | 0.300 | - | -0.799 | - | 6 | -31.313 | 3.087 | 0.037 |
| - | 0.338 | 0.253 | - | - | 0.196 | 5 | -31.260 | 3.139 | 0.036 |
| -0.162 | 0.405 | - | - | -0.528 | - | 5 | -31.075 | 3.324 | 0.033 |
| - | 0.362 | - | - | -0.369 | -0.031 | 5 | -30.119 | 4.281 | 0.020 |
| - | 0.354 | - | -0.058 | -0.339 | - | 5 | -30.116 | 4.283 | 0.020 |
| - | 0.336 | 0.302 | - | -0.499 | 0.051 | 6 | -29.931 | 4.468 | 0.019 |
| - | 0.350 | 0.291 | -0.074 | -0.573 | - | 6 | -29.891 | 4.508 | 0.018 |
| -0.145 | 0.378 | - | - | - | 0.210 | 5 | -29.613 | 4.787 | 0.016 |
| 0.023 | 0.402 | 0.157 | - | - | - | 5 | -29.480 | 4.920 | 0.015 |
| - | 0.411 | 0.153 | 0.057 | - | - | 5 | -29.478 | 4.922 | 0.015 |
| - | - | - | - | -0.577 | - | 3 | -29.454 | 4.945 | 0.015 |
| - | - | - | - | - | 0.271 | 3 | -29.163 | 5.237 | 0.013 |
| -0.198 | 0.363 | 0.282 | - | - | 0.355 | 6 | -29.053 | 5.347 | 0.012 |
| - | 0.360 | - | 0.016 | - | 0.100 | 5 | -29.035 | 5.364 | 0.012 |
| - | - | 0.264 | - | -0.767 | - | 4 | -28.880 | 5.519 | 0.011 |
| - | - | - | - | - | - | 2 | -28.635 | 5.765 | 0.010 |
| -0.262 | 0.380 | 0.346 | - | -0.643 | 0.212 | 7 | -28.566 | 5.833 | 0.009 |
| - | - | 0.268 | - | - | 0.352 | 4 | -28.518 | 5.881 | 0.009 |
| -0.002 | 0.405 | - | 0.044 | - | - | 5 | -28.506 | 5.894 | 0.009 |
| - | 0.337 | 0.255 | 0.042 | - | 0.196 | 6 | -27.854 | 6.546 | 0.007 |
| -0.193 | 0.391 | - | - | -0.461 | 0.079 | 6 | -27.783 | 6.617 | 0.006 |
| -0.162 | 0.407 | - | -0.066 | -0.540 | - | 6 | -27.687 | 6.713 | 0.006 |
| -0.186 | 0.415 | 0.302 | -0.111 | -0.824 | - | 7 | -27.612 | 6.788 | 0.006 |
| 0.194 | - | - | - | - | - | 3 | -27.511 | 6.889 | 0.006 |
| - | - | - | - | -0.366 | 0.151 | 4 | -27.270 | 7.130 | 0.005 |
| - | - | 0.312 | - | -0.497 | 0.217 | 5 | -27.176 | 7.224 | 0.005 |
| - | 0.363 | - | -0.057 | -0.379 | -0.030 | 6 | -26.721 | 7.678 | 0.004 |
| 0.033 | - | - | - | -0.534 | - | 4 | -26.673 | 7.726 | 0.004 |
| - | - | - | 0.044 | -0.570 | - | 4 | -26.649 | 7.750 | 0.004 |
| - | - | 0.130 | - | - | - | 3 | -26.570 | 7.829 | 0.003 |
| -0.067 | - | - | - | - | 0.327 | 4 | -26.424 | 7.976 | 0.003 |
| - | - | - | 0.076 | - | 0.267 | 4 | -26.378 | 8.021 | 0.003 |
| -0.145 | 0.378 | - | 0.015 | - | 0.209 | 6 | -26.194 | 8.206 | 0.003 |
| - | - | - | 0.169 | - | - | 3 | -26.190 | 8.209 | 0.003 |
| - | 0.336 | 0.302 | -0.073 | -0.515 | 0.050 | 7 | -26.173 | 8.226 | 0.003 |
| 0.024 | 0.401 | 0.159 | 0.058 | - | - | 6 | -26.086 | 8.314 | 0.003 |
| 0.033 | - | 0.265 | - | -0.725 | - | 5 | -25.821 | 8.578 | 0.002 |
| - | - | 0.265 | -0.031 | -0.773 | - | 5 | -25.789 | 8.610 | 0.002 |
| -0.118 | - | 0.279 | - | - | 0.454 | 5 | -25.683 | 8.717 | 0.002 |
| 0.219 | - | 0.167 | - | - | - | 4 | -25.549 | 8.850 | 0.002 |
| - | - | 0.269 | 0.072 | - | 0.351 | 5 | -25.451 | 8.949 | 0.002 |
| -0.198 | 0.363 | 0.283 | 0.033 | - | 0.355 | 7 | -25.261 | 9.139 | 0.002 |
| 0.191 | - | - | 0.136 | - | - | 4 | -24.786 | 9.614 | 0.001 |
| -0.264 | 0.381 | 0.348 | -0.126 | -0.674 | 0.212 | 8 | -24.448 | 9.951 | 0.001 |
| -0.110 | - | - | - | -0.399 | 0.232 | 5 | -24.385 | 10.015 | 0.001 |
| -0.171 | - | 0.329 | - | -0.554 | 0.350 | 6 | -24.372 | 10.028 | 0.001 |
| - | - | - | -0.002 | -0.366 | 0.151 | 5 | -24.174 | 10.225 | 0.001 |
| -0.193 | 0.392 | - | -0.072 | -0.473 | 0.080 | 7 | -24.020 | 10.379 | 0.001 |
| - | - | 0.124 | 0.135 | - | - | 4 | -23.836 | 10.564 | 0.001 |
| - | - | 0.313 | -0.065 | -0.510 | 0.217 | 6 | -23.777 | 10.623 | 0.001 |
| 0.032 | - | - | 0.037 | -0.529 | - | 5 | -23.585 | 10.815 | 0.001 |
| -0.068 | - | - | 0.078 | - | 0.324 | 5 | -23.359 | 11.041 | 0.001 |
| 0.219 | - | 0.167 | 0.130 | - | - | 5 | -22.537 | 11.863 | 0.000 |
| 0.033 | - | 0.266 | -0.030 | -0.731 | - | 6 | -22.405 | 11.995 | 0.000 |
| -0.118 | - | 0.279 | 0.066 | - | 0.453 | 6 | -22.285 | 12.115 | 0.000 |
| -0.110 | - | - | -0.005 | -0.400 | 0.232 | 6 | -20.964 | 13.436 | 0.000 |
| -0.174 | - | 0.332 | -0.096 | -0.575 | 0.351 | 7 | -20.617 | 13.783 | 0.000 |
